# Data-driven fine-grained region discovery in the mouse brain with transformers

**DOI:** 10.1101/2024.05.05.592608

**Authors:** Alex J. Lee, Alma Dubuc, Michael Kunst, Shenqin Yao, Nicholas Lusk, Lydia Ng, Hongkui Zeng, Bosiljka Tasic, Reza Abbasi-Asl

**Affiliations:** University of California, San Francisco; UCSF Weill Institute for Neurosciences; Allen Institute for Brain Science

## Abstract

Spatial transcriptomics offers unique opportunities to define the spatial organization of tissues and organs, such as the mouse brain. We address a key bottleneck in the analysis of organ-scale spatial transcriptomic data by establishing a workflow for self-supervised spatial domain detection that is scalable to multimillion cell datasets. This workflow uses a self-supervised framework for learning latent representations of tissue spatial domains or niches. We use a novel encoder-decoder architecture, which we named CellTransformer, to hierarchically learn higher-order tissue features from lower-level cellular and molecular statistical patterns. Coupling our representation learning workflow with minibatched GPU-accelerated clustering algorithms allows us to scale to multi-million cell MERFISH datasets where other methods cannot. CellTransformer is effective at integrating cells across tissue sections, identifying domains highly similar to ones in existing ontologies such as Allen Mouse Brain Common Coordinate Framework (CCF) while allowing discovery of hundreds of uncataloged areas with minimal loss of domain spatial coherence. CellTransformer domains recapitulate previous neuroanatomical studies of areas in the subiculum and superior colliculus, and characterize putatively uncataloged subregions in subcortical areas which currently lack subregion annotation. CellTransformer is also capable of domain discovery in whole-brain Slide-seqV2 datasets. Our workflows enable complex multi-animal analyses, achieving nearly perfect consistency of up to 100 spatial domains in a dataset of four individual mice with nine million cells across more than 200 tissue sections. CellTransformer advances the state of the art for spatial transcriptomics, by providing a performant solution for detection of fine-grained tissue domains from spatial transcriptomics data.

## Introduction

Hierarchical spatial organization is ubiquitous in tissue and organ biology. Systematic, high-dimensional phenotypic measurements of this organization, generated through experimental tools such as spatial transcriptomics, multiplex immunofluorescence, and electron microscopy, are also becoming increasingly available as large, open datasets. However, transforming this abundance of data into a useful representation can be difficult, even for fields with a wealth of prior knowledge, such as neuroanatomy.

Datasets such as the Allen Brain Cell Mouse Whole Brain (ABC-MWB) Atlas^1–3^, a multi-million cell single-cell RNA sequencing (scRNA-seq) and spatial (MERFISH) atlas, provide unprecedented opportunities to investigate whether computational tools can help biologists understand spatial cellular and molecular organization. However, the size of these datasets presents computational challenges for existing methods. Existing methods for spatial niche or spatial domain detection often operate on the entire dataset at once, for example a tissue-section-wide cell by gene matrix. This precludes scale-up to large multi-section datasets as most systems do not have the GPU memory required to load multiple sections of data or store intermediary representations such as pairwise distance matrices^4–6^, particularly as datasets scale into the millions or tens of millions. Some methods rely on Gaussian processes, which feature a costly cubic computational scaling in the number of observations^7^. Other more scalable methods are limited in capturing granular structure, integration across tissue sections, or require significant neuroanatomical prior knowledge to manually audit, cluster, and hyperparameter tune for domain discovery workflows^8,9^.

Our method, CellTransformer, implements a robust representation learning and clustering workflow to discover spatial niches at scale by representing not tissue sections but subgraphs that represent individual cellular neighborhoods. We describe an innovative strategy to induce the encoder of an encoder-decoder transformer to aggregate useful information into a neighborhood representation token. This occurs by training the model to condition cell-type specific gene expression predictions using this neighborhood context token. The model thus learns to predict expression of cell types in arbitrary cell neighborhoods. This representation allows for recovery of important anatomically plausible spatial domains while remaining computationally efficient.

We evaluate CellTransformer on using the ABC-MWB dataset (3.9 million cells collected with a 500 gene MERFISH panel)^1^ demonstrating its effectiveness in producing completely data-driven spatial domains of the mouse brain by comparing the results to the Allen Mouse Brain Common Coordinate Framework version 3 (CCFv3)^10^. CCF is a consensus hand-drawn 3D reference space compiled from a large multimodal data corpus. Annotations feature labels at three levels of coarseness (from 25 regions at coarse-grain to 670 at fine-grain), which we use to show that CellTransformer excels at identifying spatial domains which are spatially coherent and biologically relevant. CellTransformer domains reproduce known regional architecture observed in targeted studies of the subiculum and in the superior colliculus superficial layers. Beyond the 670 regions currently annotated in ABC-MWB, we show our workflow produces meaningful data-driven domains in regions which currently lack subregion annotation. As examples, we focus on data-driven subdomains we define in superior colliculus and midbrain reticular nucleus.

We also demonstrate CellTransformer’s strength in integrating domains across animals, leveraging a separate whole-brain dataset within ABC-MWB^11^ comprising 6.5 million cells distributed across four animals and 239 sections and with a separate gene panel with 1129 genes. We find that CellTransformer produces consistent subregions across all 5 animals (1 coronal and 4 sagittal), suggesting a successful integration across animals with heterogeneous measurements. Notably we also find that identified domains are highly consistent across animals. To our knowledge, this work provides the first demonstration that large scale data-driven discovery of domains at CCF-like resolution can be based on spatial transcriptomics data. Finally, we show that our framework can perform domain detection in a different spatial transcriptomics modality, Slide-seqV2, using the whole-brain dataset of cellularly deconvoluted results^12^.

## Results

### The CellTransformer architecture and domain detection workflow

CellTransformer is a graph transformer^13^ neural network that is trained to learn latent representations of cell neighborhoods by conditioning single-cell gene expression predictions on neighborhood spatial context. We define a cellular neighborhood as any cells within a user-specified distance cutoff in microns away from a reference or center cell. As input, our model requires the gene expression profiles and cell type classifications for cells in a neighborhood and outputs a latent variable representation for that neighborhood. One of the principal operations in a transformer is the self-attention operation, which computes a feature update based on pairwise interactions between elements in a sequence, which are referred to as tokens (here, cells). Accordingly, one interpretation of our model is of learning an arbitrary and dynamic pairwise interaction graph among cells.

Restricting this graph to a small neighborhood subgraph of the whole-tissue-section graph has benefits for both computational resource usage and biological interpretability. We interpret the size of the neighborhood as a constraint on the physical distance at which statistical correlations between the observed cells and their gene expression profiles can be directly captured. Truncating neighborhoods using a fixed spatial threshold instead of choosing a fixed number of neighbors also allows the network to account for the varying density of cells in space. Accordingly, our framework incorporates a notion of both cytoarchitecture (relative density and proximity) and molecular variation (cell type and RNA-level variation) in the data.

To induce our model to learn biologically relevant latent features from cell neighborhoods, we designed a self-supervised training scheme requiring only cell-type labels, which many large-scale studies make available via scRNA-seq atlas reference mapping^1,11^. Specifically, we train the model to extract features from cellular neighborhoods, modeled as sets of cell tokens that are within a box of fixed size centered around a center, or reference cell, and use them to predict the observed gene expression of the cell at the center of the neighborhood. We refer to this cell as the reference cell (indicated by “cell R” in **Figure 1a**). Cell tokens are generated by composing cell-type and gene expression information (**Methods**). After encoding with a series of transformer layers (where cells are only allowed to attend to each other if they are in the same neighborhood), these tokens are then aggregated using a learned pooling operation to produce a single token representation of the entire tissue context. The model receives a new mask token representing the reference cell’s type which is used to predict its gene expression following the operation of several transformer decoder layers (**Figure 1b**). Importantly, during this process, only the mask token and the neighborhood representation can attend to each other. This operation captures a hierarchical encoding and decoding process where low level information (gene and cell type) is produced at the cell token level and aggregated into a high-level representation. This high-level representation is then used to conduct the reverse decoding process (prediction of gene expression from cell type and tissue context information). Unlike closely related method NCEM^14^, which predicts expression of a reference masked node, we aggregate information across tokens (nodes) in a cellular neighborhood using a learned pooling which strongly bottlenecks the information distributed across the tokens prior to masked cell prediction.

**Figure 1.**
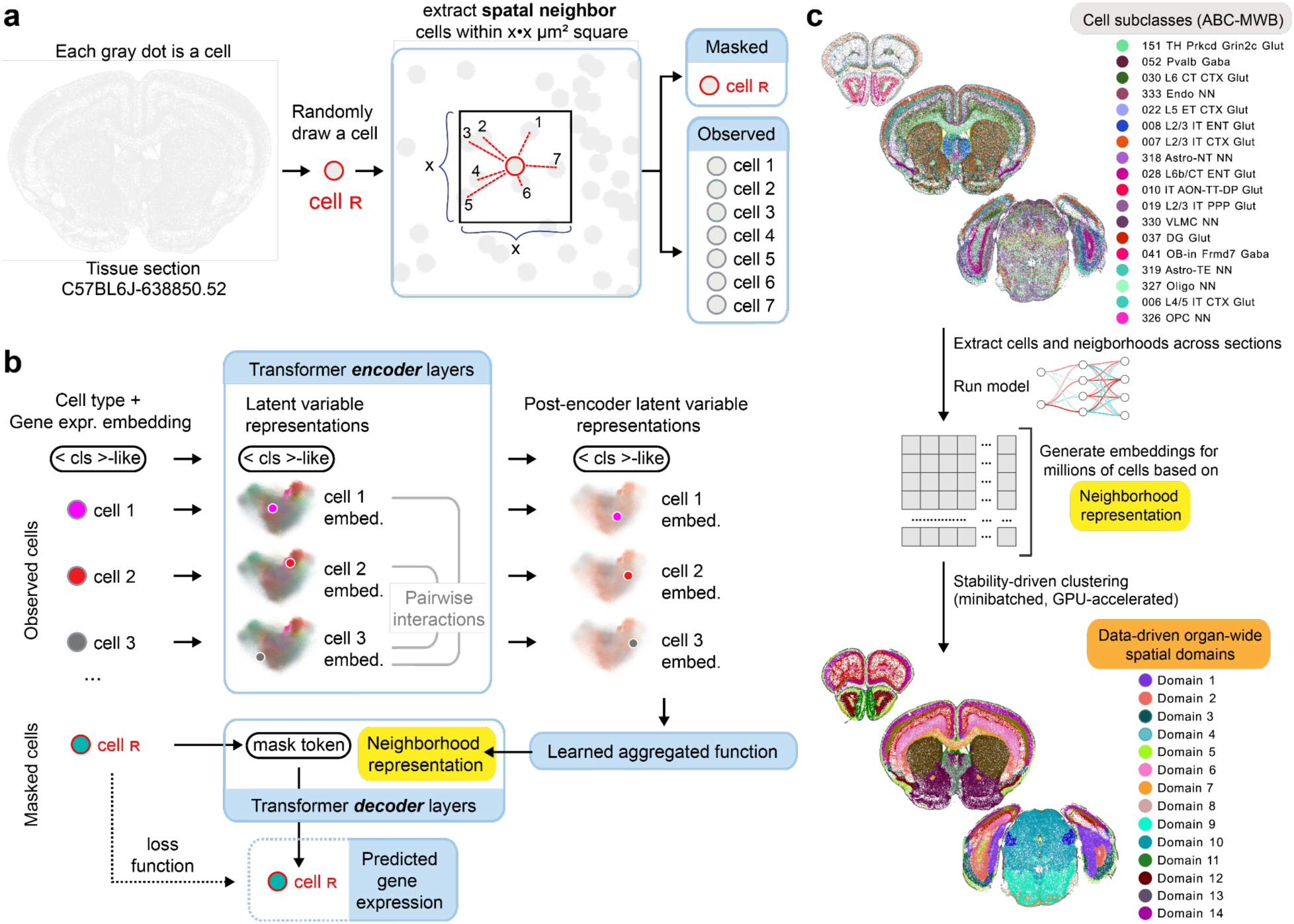
Overall training and architectural scheme for CellTransformer. (**a.**) During training, a single cell is drawn (we denote this the reference cell, highlighted in red). We extract the reference cell’s spatial neighbors and partition the group into a masked reference cell and its observed spatial neighbors. (**b.**) Initially, the model encoder receives information about each cell and projects those features to *d-*dimensional latent variable space. Features interact across cells (tokens) through the self-attention mechanism. These per-cell representations and an extra token acting as a register token are then aggregated into a single vector representation, which we refer to as the neighborhood representation. This representation is concatenated to a mask token which is cell type-specific and chosen to represent the type of the reference cell. A shallow transformer decoder (dotted lines) further refines these representations and then a linear projection is used to output parameters of a negative binomial distribution modeling of the MERFISH probe counts for the reference cell. (**c.**) Once the model is trained, we compute embeddings (one for each neighborhood/reference-cell pairing) and concatenate these embeddings within the tissue section datasets and across tissue sections. Concatenating embeddings across tissue sections produces region discovery at organ level. We then cluster these embeddings using *k-*means to discover tissue domains across sections.

At test time, we extract this neighborhood representation for each cell and use *k-*means clustering to identify discrete spatial domains (**Figure 1c**). We will use the term spatial domain to refer to the output of clustering on embeddings and cluster to refer to single-cell clusters transferred from the ABC-WMB single cell taxonomy. We emphasize that the input embedding matrix for *k-*means is conducted by concatenating all cells across the dataset across tissue sections. Since minibatching is used during training (unlike methods such as STAligner and GraphST), for generating embeddings, and during *k-* means (using cuml for GPU-acceleration), overall computational costs of our algorithm are limited in principle only by the memory required for storage of cellular neighborhoods rather than entire sections or datasets.

### Data-driven discovery of fine-grained spatial domains in the mouse brain using ABC-WMB

The ABC-WMB spatial transcriptomics dataset contains data from five mouse brains^1,11^. One animal was processed by the Allen Institute for Brain Science with a 500 gene MERFISH panel and 53 coronal sections (Yao et al, 2023)^1^ The remaining four other animals, generated in Zhang et al. (2023)[11] were collected with a 1129 gene panel. Sections from two of these animals (“Zhuang 1”, 147 sections; and “Zhuang 2”, 66 sections) were sampled coronally. The other two animals in the dataset (“Zhuang 3”, 23 sections; and “Zhuang 4”, 3 sections) were sampled sagittally.

We first trained CellTransformer on the Allen 1 dataset, subsequently extracting embeddings for each cell’s neighborhood, which we defined as a set of cells within a fixed size square around that cell. We then clustered these embeddings using *k-*means. We emphasize that to generate spatial domains across the brain, all *k-*means clustering in this paper was performed by concatenating cells in the dataset across tissue sections. All further references to visualizations of domains, including those only visualized for a subset of domains, were fit at a given number of domains across the entire dataset. We also optionally introduced a smoothing step prior to *k-*means, which we applied to spatially smooth the embeddings. See **Supplementary Note 1** for a discussion on the effects of smoothing on detected domains.

We generated domains at *k=*25, 354, and 670, to match the division, structure, and substructure annotations in CCFv3, displaying domains for four consecutive tissue sections (**Figure 2a**). We also provide representative images of spatial clusters across the brain (28/53 sections) at different *k* in **Supplementary Figures 1-3**. Low domain numbers such as *k*=25 broadly divide the brain into neuroanatomically plausible patterns, with subregions of striatum (dorsal and ventral marked in **Figure 2a**) and cortical layers clearly visible. A comparison of cortical layers across these sections shows that CellTransformer domains at *k=25* are well matched to CCF (**Supplementary Figure 4b**) and correctly identify major classes of layers (1, 2/3 4, 5, and 6) across somatosensory and somatomotor cortex. In particular, we point out the excellent correspondence of domains across tissue sections at *k*=25 across the entire dataset (**Supplementary Figure 1**), with nearly perfect consistency across regions. This suggested that our neighborhood representation method was robust enough to enable integration without modeling of batch or tissue-level covariates.

**Figure 2.**
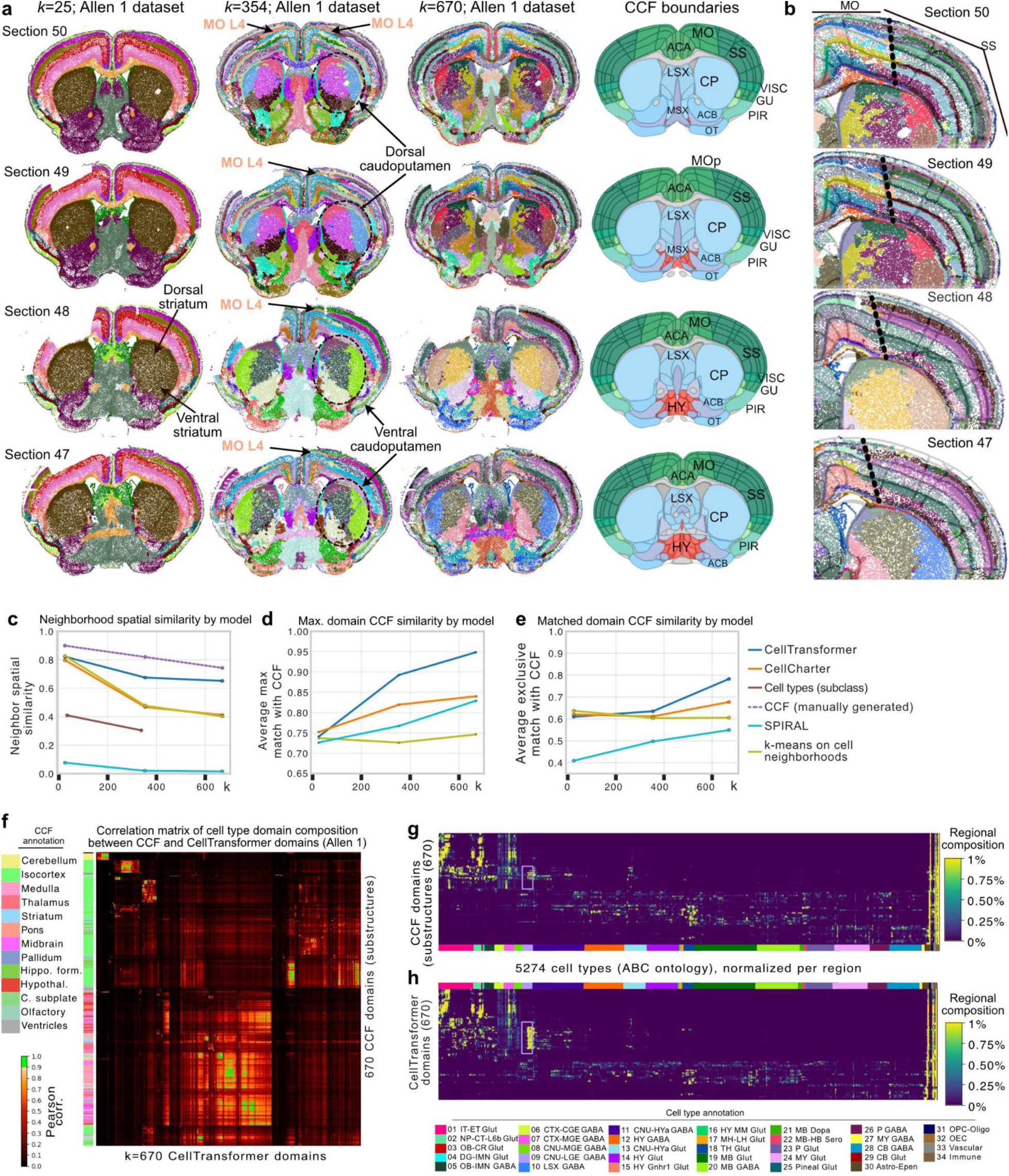
Representative images of spatial domains discovered using CellTransformer on the Allen 1 dataset (53 coronal sections and 500 gene MERFISH panel^1^) and comparison to CCF. (**a**.) Four sequential tissue sections (the inter-section distance is 200 µm) from anterior (first row, corresponding to section 50) to posterior (bottom row, section 47). In the first three columns, each dot is a cell, colored by spatial domain identified by CellTransformer when clustering was conducted with *k* = 25, 354, and 670 domains (the CCF division, structure, and substructure domain resolutions). Spatial domain labels are depicted with the same colors across sections within the same column. Fourth column shows CCF region registration to the same tissue section. Select regions are annotated with CCF labels. MO: motor cortex, SS: somatosensory cortex; ACA: anterior cingulate, CP: caudoputamen; LSX: lateral septum; MSX: medial septum; VISC: visceral cortex; GU: gustatory cortex; PIR: piriform cortex; OT: olfactory tubule; ACB: nucleus accumbens; HY: hypothalamus (**b.**) Single hemisphere images of same tissue sections in **a.** domains fit at *k*=670, zoomed in on cortical layers of motor cortex (MO) and somatosensory cortex (SS). CCF boundaries are shown in semi-transparent lines, with the boundary between SS and MO outlined in larger black dotted lines. (**c.**) Spatial homogeneity (see **Methods**) of domains from different methods including recently published methods CellCharter and SPIRAL. (**d.**) Average Pearson correlation (averaging over number of domains and method) of the maximum Pearson correlation between the cell type composition (at subclass level, 338 types) vectors of data-driven regions with CCF ones. (**e.**) Average Pearson correlation (averaging over number of domains and method) of optimal matched pairs between data-driven and CCF regions, where CCF regions are only allowed to pair with one data-driven region per comparison. Matches fit using linear programming. (**f.**) Region-by-region Pearson correlation matrix comparing cell type composition vectors from 670 CCF regions (at substructure level) with 670 spatial domains from CellTransformer. The CCF regions are shown on the left with their structure annotations from CCF at division level on the side of the plot. Correlations above 0.9 are shown in bright green to assist in visualization. (**g.**) Cell type (cluster level) by region matrix for 670 CCF regions at substructure level. (**h.**) Cell type (cluster level) by region matrix for 670 CellTransformer regions. Rows are normalized to sum to 1 in both **g.** and **h.** Colors along x-axis in both **g.** and **h.** show cell class annotations from ABC-MWB cell type taxonomy at class level to allow for visualization of composition in terms of known types. Cell types in the “09 CNU-LGE GABA” class are boxed in purple in **g.** and **h.**, matching their color in the legend. Rows of both **g.** and **h.** are grouped using clustering to produce approximately similar structure.

At *k*=354, anterior-posterior subdivisions emerge such as the presence of layer 4 in the motor cortex^13^ (**Figure 2a**, see **Supplementary Figure 4d, e**). Historically, the mouse motor cortex was thought to lack a granular layer 4, however recently, MERFISH, transcriptomic and epigenomic studies have confirmed its existence^1,15,16^. At *k=*100 and *k=*354, we find a domain corresponding to Layer 4 in the somatosensory cortex which clearly extends to layer 4 in the motor cortex.

At *k=*670, the cortical layers identified at lower resolution are further partitioned into superficial, intermediate, and deep strata within several layers. We visualize cortical layers across sections in depth (**Figure 2b**), showing CellTransformer not only identifies fine superficial-deep structure within cortical layers but also preserves the boundary between somatosensory and motor cortex (marked in thick black dotted lines in **Figure 2b**). Taken together these results showed that CellTransformer robustly describes previously known anatomical structures.

We also examined the caudoputamen at various choices of *k.* At *k*=25, the caudoputamen is one domain, which separates into broad spatially contiguous domains at *k*=100. Interestingly, at *k*=354 and *k*=670, we observe domains that intermingle in a grid-like pattern (**Figure 2a**, **Supplementary Figure 5**) that strongly resembles the Voronoi parcellation established in Hintiryan et al. (2016)^17^ through systematic projection mapping to caudoputamen. Notably, CellTransformer also captures the transition between the quadrant pattern in intermediate caudoputamen (sections 52, 50 and 49 in **Supplementary Figure 5**) to the sequential strip organization (sections 44, 43) which Hintiryan et al. (2016) attributed to the differences in subnetwork reorganization. The correspondence of our transcriptomic domains to the Hintiryan et al. (2016) results, which are exclusively based on projection mapping (non-transcriptomic data), suggests the biological relevance of our representation learning workflow.

We compared CellTransformer to several other workflows to capture spatial coherency and multiresolution neuroanatomical annotations in CCF at the division, structure, and substructure levels. For comparison, we used two recent methods, CellCharter^18^ and SPIRAL^19^ that are scalable to millions of cells as benchmarks. CellCharter builds spatially informed embeddings for domain detection by concatenating the embeddings across scales followed by dimensionality reduction and batch correction, while SPIRAL uses graph-neural networks for batch effect correction and integration across scales. Additionally, we implemented a machine learning baseline that employs *k-*means clustering on cellular neighborhoods (represented as cell type count vectors). Many of the other GPU-accelerated methods such as scENVI^4^, STACI^20^, spaGCN^5^, STAligner^6^, STAGATE^21^ or GraphST^22^ cannot be run on datasets that contain millions of cells due to computational constraints (see **Methods**). Many of these methods require instantiation of a dataset-wide pairwise distance matrix between all cells either on GPU or in RAM, which is a prohibitively large matrix (∼60TB for ∼4M cells) even for enterprise-level hardware. In contrast, our workflow does not require very large system RAM or extensive preprocessing steps due to our training and inference, maxing out at less than 100GB but requiring significantly less in practice.

To quantify the spatial coherence of domains, for each cell we identified its nearest 100 spatial neighbor cells. We then quantified the proportion of neighbor cells within the same spatial domain label as the starting cell (**Figure 2c**). Ideally, we would expect a high proportion of neighbor cells to be in the same spatial domain as the starting cell. In this comparison of neighborhood spatial smoothness, CellTransformer outperforms CellCharter (58.2% better spatial coherence at 670 domains) and SPIRAL (4091.2%). CellTransformer also outperforms the machine learning baseline based on *k-*means clustering on cellular neighborhoods (61.9% better spatial coherence). For reference, we include the CCF parcellation (dashed purple line) in this comparison to provide an upper bound, as well as spatial coherence using single cell type calls at subclass level (338 types, see **Methods**).

To quantify the similarity of detected domains with CCF annotations, we compared the cell type composition of domains using cell type calls from the ABC-WMB taxonomy. We again chose the subclass cell type level, extracting for each domain and for each method a 338-long cell-type vector. We calculated the Pearson correlation of cell type composition vectors computed using the CCF regional annotations at division (25), structure (354) and substructure (670) levels against those of the various methods at the corresponding number of spatial domains. First, for each data-driven domain, we computed the maximum correlation to any CCF domain at the same CCF annotation resolution averaging these numbers across domains. CellTransformer outperforms other methods at mid-granularity and fine-granularity (**Figure 2d**). In this comparison, several data-driven regions can match the same CCF region, which in the worst case could provide an overly optimistic picture of the correspondence between data-driven domains and CCF. To address this, we conducted a second analysis where only one CCF region could be matched to a given data-driven one. We used linear programming to optimize 1:1 pairing of data-driven regions to CCF ones based on their Pearson correlation and averaged these across regions and methods (**Figure 2e**).

CellTransformer is highly performant, showing that increase in correlation is not due to redundant matches to a single area in CCF. Visualization of spatial clusters from CellCharter (**Supplementary Figure 6-7**) at *k=*670 domains across the brain and in midbrain shows lack of spatial coherence in cortical layers and midbrain, with detected domains distributed in a what appear to be non-biological patterns. In contrast, CellTransformer identified spatially coherent domains and uncovered plausible neuroanatomical structures.

To further characterize the similarity of CellTransformer domains with CCF, we plotted the Pearson correlation matrix (**Figure 2f**) between cell type composition vectors generated at 670 domains (substructure level in CCF). Block structures with very high correlations (>0.9, shown in bright green) in the matrix clearly show that CellTransformer is able to identify regions that are highly similar with known ones without any labels. We also investigated correspondence of cell type composition with more granular single cell annotations, employing the “cluster” (5274 cell types) level annotations from ABC-MWB. We observed high similarity between the “substructure” CCF domain set (**Figure 2g**) and 670 CellTransformer domains (**Figure 2h**) with average Pearson correlation of CellTransformer to CCF domains of 0.853. This shows the high correspondence of CCF and CellTransformer (**Figure 2g** and **Figure 2h**) is robust to cell type resolution at which comparison occurs. CellTransformer identified an increase in number of domains containing the 09 CNU-LGE GABA class (striatal/pallidal GABAergic neurons from lateral ganglionic eminence compared with the 670 CCF substructures, shown in light purple box in **Figure 2g** and **Figure 2h**), potentially suggesting the presence of uncharacterized developmental populations.

The observation of hierarchical grouping of domains at different choices of *k* (for example delineation of cortical layers and sublayers with increasing number of domains) prompted us to develop a strategy to evaluate an optimal number of spatial domains based on two metrics. We implemented a previously published strategy^23^ to determine the optimal number of domains using a stability criterion. We reasoned that the optimal choice of spatial domain number would feature minimal variability across clustering runs. In brief, we computed 20 clustering instances with different random initializations for a large range *k* values (100-2000) and quantified their variability over these initializations (see **Methods**). Interestingly, stability increased with increasing *k* (**Supplementary Figure 8a, 8b**). To facilitate the choice of a particular resolution for analysis, we also computed the inertia (sum of squared errors) for each clustering solution. Low stability at small numbers of domains may partially explain subpar results for CellTransformer in the *k*=25 CCF evaluations. We averaged the inertia curve and instability and computed the point of second derivative crossing to identify *k=*1300 as our resolution for analysis (crossing point shown with red dot in **Supplementary Figure 8c**).

CellTransformer is the only method out of the three we implemented (including six other pipelines which were unable to cope with the size of ABC-MWB dataset) to allow discovery of spatially coherent divisions at greater than CCF resolution. To our knowledge, this study establishes the first instance of a data-driven method using spatial transcriptomics data to identify brain regions at resolutions finer than previously defined in the CCF. We next sought to establish correspondence of particular domains at *k*=1300 to known neuroanatomy.

### Mapping of spatial domains in the hippocampal formation

We characterized CellTransformer’s ability to capture known anatomical structure in the hippocampal formation, notably the subiculum (SUB) and prosubiculum (PS), in the Allen 1 dataset. We focused on this area because it is well characterized with respect to both connectivity^24^ and transcriptomic composition^25,26^. These structures were investigated in Ding et al. (2020)^27^, where the authors performed consensus clustering of glutamatergic neurons and subsequent ISH experiments were used to comprehensively map domains in dorsal subiculum (SUBd) and dorsal and ventral prosubiculum (PSd and PSv). Specifically, this and other recent works have noted the extensive laminar organization (superficial layers to deeper layers), and the dorsal-ventral organization of the subiculum^28–30^. This organization has been attributed to distinct and correlated patterns of gene expression and connectivity.

We qualitatively compared spatial domains discovered by CellTransformer with *k=*1300 to the anatomical borders identified in Ding et al. (**Figure 3a**). The subiculum features a three-layer organization referred to as molecular (mo) layer, a pyramidal cell (py) layer, and polymorphic (po) cell layer. **Figure 3a** shows a diagram of SUB and PS regions based on Ding et al. (2020) with the pyramidal and polymorphic layers of SUB and PS annotated in bold black text. **Figure 3b** shows discovered spatial domains at *k*=1300 across four sequential sections corresponding to those in Ding et al. (2020). A subset of domains corresponding to SUB and PS are shown in **Figure 3c** along with putative regional annotations.

**Figure 3.**
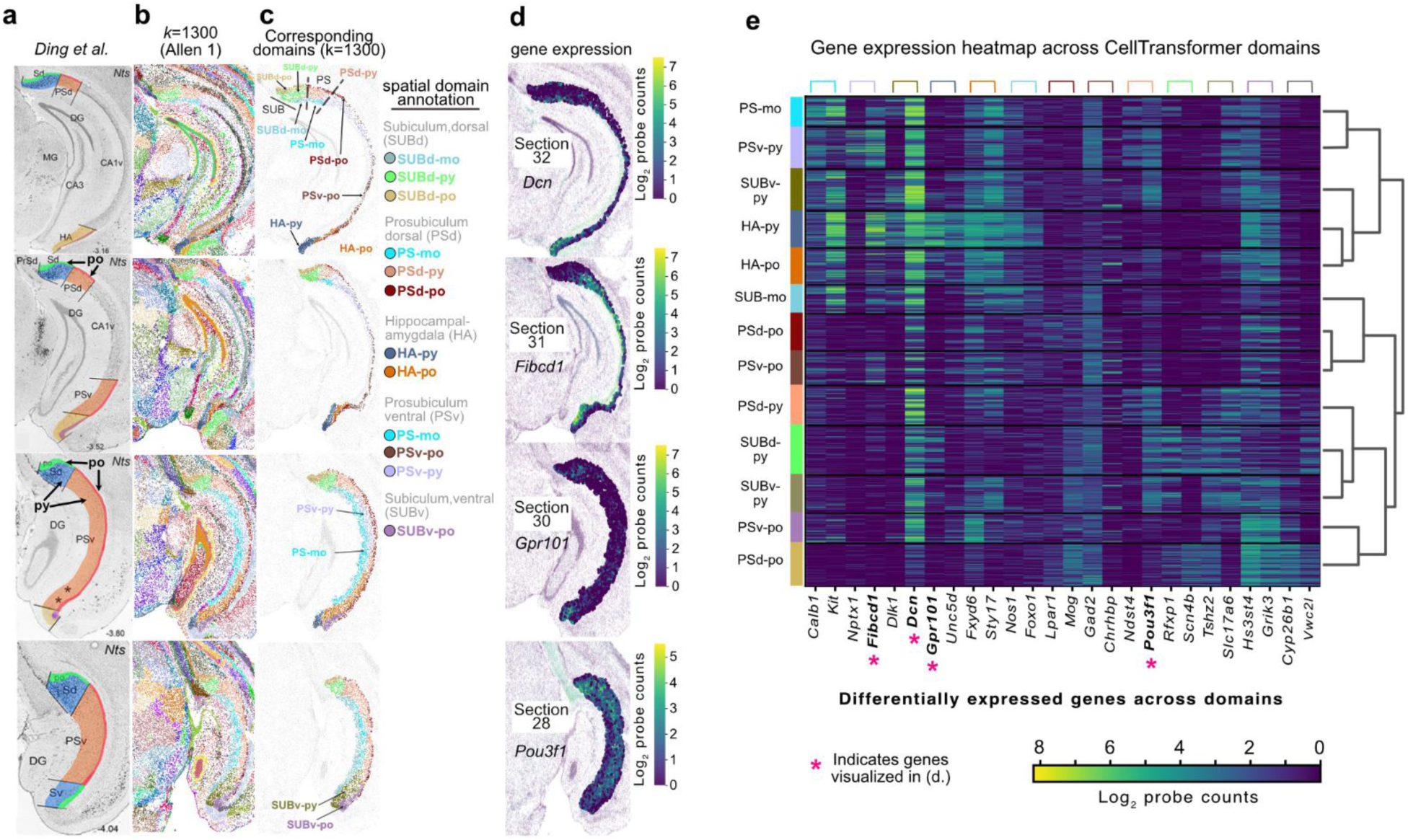
Comparison of CellTransformer domain sets identified in the Allen 1^1^ dataset with *k=*1300 with a comprehensive region set found in Ding et al. (2020), reproduced with permission from authors. (**a.**) Representative images reproduced from Ding et al. of region boundaries in prosubiculum (PS), subiculum (SUB), and hippocampal-amygdala (HA) particularly along the dorsal-ventral axis. Polymorphic and pyramidal layers of the dorsal subiculum (SUBd) and ventral prosubiclum are indicated. (**b.**) Images from hippocampal formation across 4 sequential tissue sections (anterior to posterior) roughly aligned to sections presented in Ding et al. Each dot is a cell colored using domain labels with *k=*1300. (**c.**) Same as **b.** but only showing cells inside PS, SUB, and HA. Putative regional annotations are indicated and grouped by dorsal or ventral region within PS and SUB. (**d.**) Gene expression patterns visualized at the corresponding tissue section, where only cells within PS/SUB/HA are shown. Units are in log_2_ probe counts. (**e.**) Gene expression heatmap of identified subregions, with putative anatomical annotation. Dendrogram from hierarchical clustering in gene expression space is shown to the right. Genes visualized in **d.** are bolded and denoted with a pink asterisk. Colored brackets indicate the genes which are differentially expressed with respect to the domain (colors match those shown on the left of the heatmap). Two genes per domain are shown and each gene is expressed with at least log-fold change greater than 1 relative to the other domains. Abbreviations: PS-mo: prosubiculum molecular layer; PS-py; pyramidal layer of subiculum; SUBv-py; ventral subiculum, pyramidal layer; HA-py: hippocampal-amygdaloid transition area, pyramidal layer; HA-po: hippocampal-amygdaloid transition area, polymorphic layer; SUB-mo: subiculum, molecular layer; PSd-po: dorsal prosubiculum, polymorphic layer; PSv-po: ventral prosubiculum, polymorphic layer; PSd-py: dorsal prosubiculum, pyramidal layer; SUBd-py: dorsal subiculum, pyramidal layer; SUBv-py: ventral subiculum, pyramidal layer; PSv-po: ventral prosubiculum, polymorphic layer; PSd-po: dorsal prosubiculum, polymorphic layer.

CellTransformer identifies a three-layer organization in the dorsal subiculum corresponding to that in Ding et al. (2020) labeled SUBd-py (light green), SUBd-po (gold), and SUBd-mo (gray-blue). CellTransformer also correctly splits the SUBd and PSd shown with black dotted lines on the image of section 32. Three-layer strata are also observed in PSd, although notably the pyramidal layer domain extends caudally, consistent with transcriptomic studies^24–26^ of SUB architecture. For instance, our PSd-po domains (sections 31 and 30) strongly resemble the HGEA layer 4 found in Bienkowski et al. (2018)^24^. Note that differences may arise between panels in **Figure 3a** and **3c** because of sectioning variability and lack of exact match between sections in ABC-MWB and the Ding et al. (2020) study. In addition to the aforementioned regions we also observe high agreement in areas such as in the hippocampus-amygdaloid transition area (HA) and ventral prosubiculum (PSv).

Ding et al. (2020) observed differential projection topology in dorsal subiculum versus ventral prosubiculum. Correspondingly, genes were found to form opposing gradients across the length of subicular areas. Dorsolateral gene gradients appeared in SUBd and ventromedial gradients in PSv. Since CellTransformer domains appeared to correspond well with literature results, we explored gene expression patterns across domains to verify whether dorsal-ventral and medial-laterally varying gene patterns could be observed. We conducted differential expression analysis across our subicular domains (**Figure 3e**) that when visualized (**Figure 3d**) clearly reflected these gradients. Many genes expressed in SUB and PS traverse their long axis as reported previously^25^. The identification of spatial domains which subdivided specific layers of PS and SUB similarly to the results in Ding et al., and featured similar types of gene expression gradients as existing literature, suggests that our pipeline was successful in learning neuroanatomically useful information. Importantly, while results in Ding et al. and related works were enabled by significant neuroanatomical and experimental expertise, CellTransformer allows identification of granular tissue structure in a data-driven fashion. Encouraged by this result, we continued our investigation of CellTransformer correspondence with known literature with a comparison in superior colliculus.

### CellTransformer allows for quantification of laminar and columnar organization in superior colliculus

Recent studies using systematic mapping of cortico-tectal fibers in superior colliculus (SC) have identified distinct laminar and columnar structure^31^, suggestive of the complex role SC plays in integration of sensory information and coordination of signals. Therefore, SC presented an excellent opportunity to identify transcriptomic and cellular correlates of connectomic variation. We observed a strong correspondence of three of our spatial clusters (*k*=1300) in the Allen 1 dataset with known layers of superior colliculus, sensory area, particularly the zonal (zo), superficial gray (sg), and optic (op) layers across a set of tissue sections spanning ∼600 µm from anterior to posterior (rows of **Figure 4a** and **Supplementary Figure 9a**). CellCharter was unable to identify these structures (**Supplementary Figure 7)** and only identifies two layers in SC, which does not conform with existing results.

**Figure 4.**
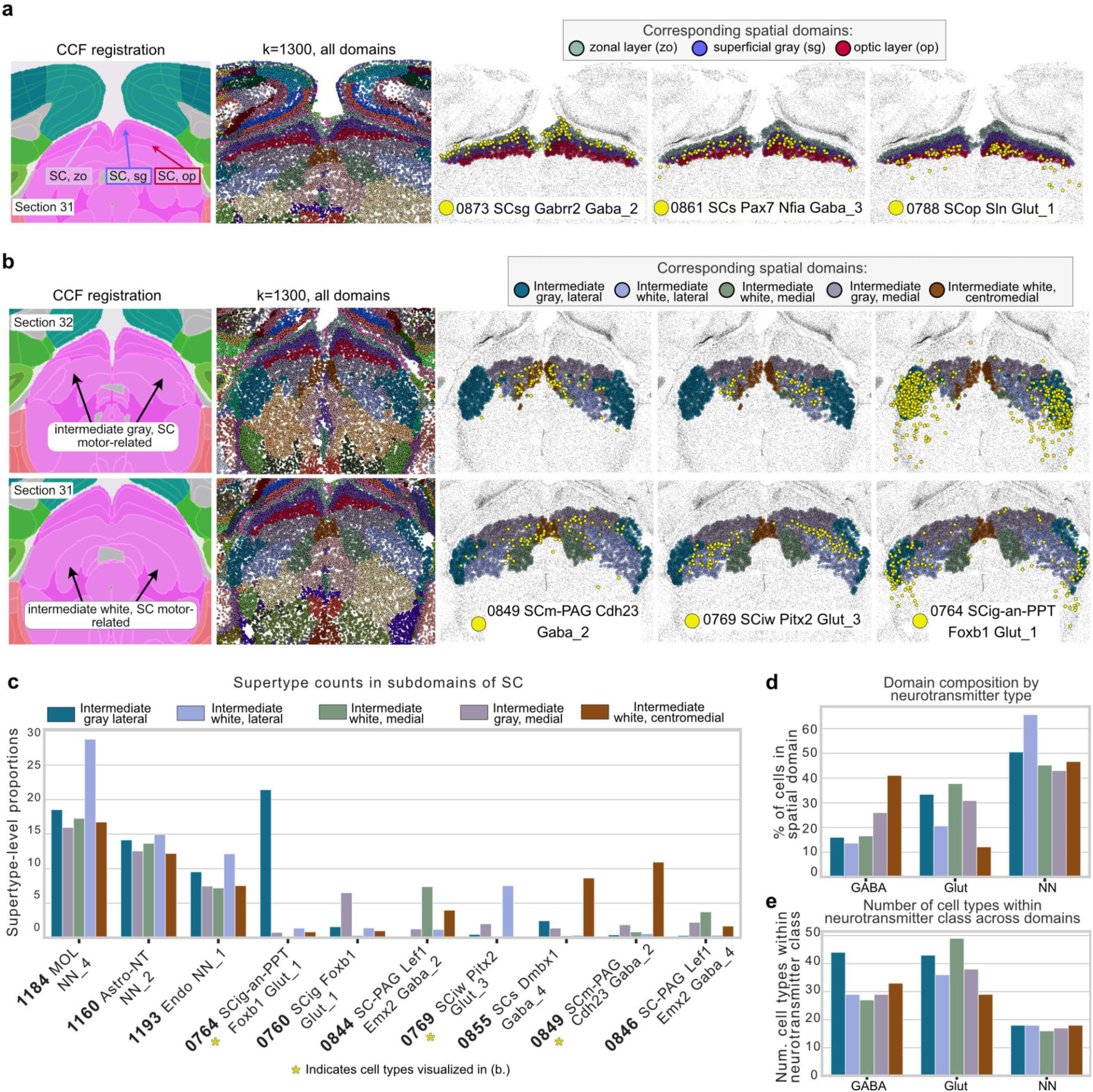
Examination of putative dorsal and ventral subregions of superior colliculus identified in the Allen 1^1^ dataset using CellTransformer. (**a.**) Putative subregions of sensory layers of superior colliculus in tissue section 32 identified with *k*=1300 CellTransformer domains. CCF registration is in the first column, with zonal (zo), superficial gray (sg), and optic (op) layers labeled by the color of their CellTransformer domain in third, fourth, and fifth columns. The second column shows all cells with color labels from their spatial domain from CellTransformer at *k*=1300. The third, fourth, and fifth columns show the putative zo (gray-green), sg (purple), and op (red) domains. These columns also show the spatial distribution of one supertype level cell type in yellow across the section. (**b.**) Sequential tissue sections (32: anterior, 31: posterior) shown similarly to **a.**, but visualizing subregions of the intermediate gray and intermediate white layers, which are indicated with black arrows in the CCF registered annotation image. (**c.**) Proportions of different supertype level cell types for top-ten most abundant types in different spatial domains. Colors refer to the same spatial domain label in (**a.**) and (**b.**). Cell types visualized in (**b.**) are denoted with a yellow asterisk. (**d.**) Barplot of the percentage of cells of a given neurotransmitter class found in a given region (GABA - GABAergic; Glut - Glutamatergic; NN - non-neuronal). (**e.**) Number of unique cell types (at supertype level) found in each domain, grouped by neurotransmitter class.

By visualizing the cell type composition within the top-ten most abundant types for these three spatial domains (**Supplementary Figure 9a, 9b**), we were able to identify cell types that were highly selective for our data-driven SC layers: types 0873 SCsg Gabrr2 Gaba_2, 0861 SCs Pax7 Nfia Gaba_3, and 0788 SCop Sln Glut_1. Crucially, the cell types, which have already been annotated as being associated with one of the zonal, optic, or superficial gray, are identified automatically by CellTransformer. We chose the supertype level to allow inspection of abundant cell types without being difficult to visualize. Supertype-level visualizations also show that even with granular cell types (1201 types in Yao et al.) CellTransformer domains are often marked by spatially specific cell type patterning; we note that we do not filter cells outside of our putative superior colliculus layers for visualization. Next we visualized the percentage of cells in each domain (**Supplementary Figure 9c**), grouping them by neurotransmitter class (GABA-ergic, glutamatergic, and non-neuronal). The superficial gray layer showed the higher proportion of GABA-ergic neurons, while the optic layer had the highest proportion of glutamatergic neurons. To further explore these relationships, we calculated the number of distinct cell types (supertype level) within each neurotransmitter class and domain. A clear dorsal-ventral organization was evident (**Supplementary Figure 9d**) with the number of GABA-ergic and glutamatergic neuron types increasing with layer depth, suggesting CellTransformer’s ability in capturing complex patterns of cellular spatial organization.

Encouraged by these findings, we also investigated subregions of the intermediate gray and intermediate white areas of the motor-related areas in SC (**Figure 4a-b**), where we identify consistent regions across two consecutive sections that are not annotated in the CCF (rows of **Figure 4b**). We define subregions of intermediate gray (ig) and white (iw), noting a medial-lateral structure similar to that in Benavidez et al. (2021)^31^, which exhaustively cataloged projection zones in superior colliculus. Notably unlike in superior colliculus sensory, a significant number of non-neuronal cell types are found in very similar proportions across the intermediate white and gray layers (**Figure 4c**), and instead differences in regions may be attributable to varying proportions of rare cell types. Encouragingly, even in these fine-grained areas, cell types that are highly specific for our data-driven layers can be readily identified (columns of **Figure 4b**). These rare domain-enriched cell types include: 0849 SCm-PAG Cdh23 Gaba_2 (enriched in the medial intermediate white layer, shown in dark green), 0769 SCig SCiw Pitx2 Glut_3 (enriched in lateral intermediate white, shown in light blue), and 0764 SCig-an-PTT Foxb1 Glut_1 (enriched in medial intermediate gray, shown in dark blue). The identification of *Pitx2*-expressing neurons also supports our assertions that CellTransformer identifies biologically relevant domains, with previous studies using *Pitx2* expression specifically as an intermediate layer marker in superior colliculus^16,28^.

We observed complex cell type abundance gradients when visualizing the percentage of cells in a given domain by their neurotransmitter type (**Figure 4**). We used supertype level to confirm that spatially-varying cell distribution patterns persisted when using more granular cell type annotations. Lateral domains such as intermediate gray, lateral (shown in dark blue) and intermediate white, lateral (shown in light blue) featured a smaller proportion of GABA-ergic neurons than medial domains but were enriched for glutamatergic neurons and non-neurons (**Figure 4d**). Despite the low proportion of GABA-ergic neurons, the lateral domain of intermediate gray possessed the highest number of GABA-ergic neuron cell types (**Figure 4e**). Conversely, non-neurons featured the same number of types across the laminae.

### A medial-lateral gradient of inhibitory neurons in the midbrain reticular nucleus

Next, we investigated the midbrain reticular nucleus (MRN), a subcortical structure with few anatomical annotations in CCF. MRN is highly enriched for interneurons in a dense array of connections and appears to play a complex role in movement initiation and release^29,30^. CellTransformer identifies four subregions of the MRN, which are not included in the existing CCF annotation (**Figure 5a**). Plotting cell type proportions across the MRN, we identified cell types which are enriched for these putatively uncharacterized areas, although all domains were predominantly glial (e.g., 1184 MOL NN_4 supertype is abundant in all regions, **Figure 5b**). By visualizing differentially expressed genes across the domains (**Figure 5c**), we identified genes which were subregion selective (selected genes shown in **Figure 5d**) and form dorsal-ventral expression gradients. Hierarchical clustering showed that the two dorsal domains (purple and brown) group together with the two ventral ones (gold and gray).

**Figure 5.**
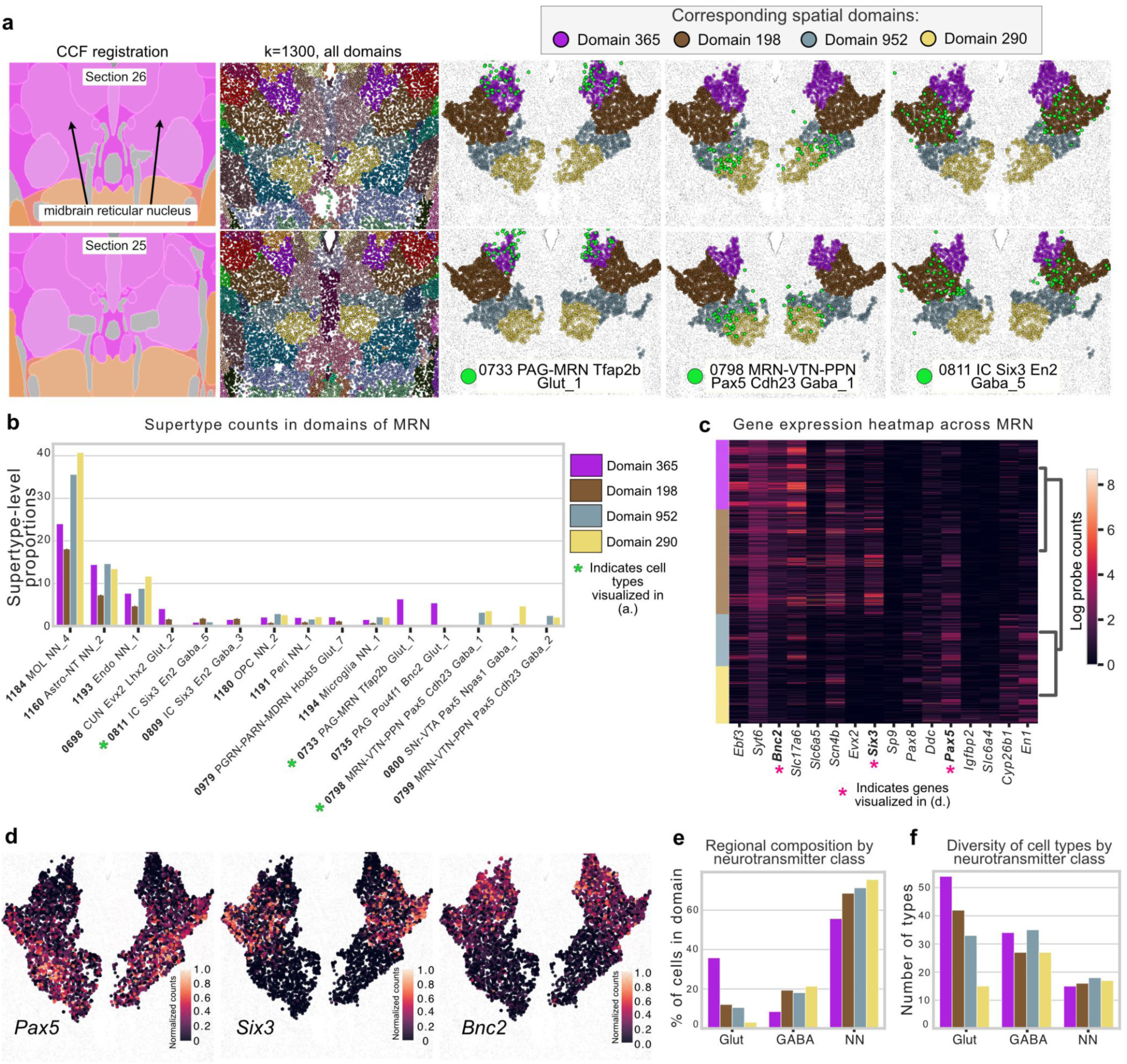
Subregions of midbrain reticular nucleus discovered in the Allen 1^1^ dataset using CellTransformer. (**a.**) Sequential tissue sections (26 and 25, anterior to posterior). First column: CCF registration with approximate location of midbrain reticular nucleus shown with arrows. Note that registration is not exact and can differ across hemispheres. Second column: all cells in field of view, with color from spatial domains determined with *k*=1300. The rest of the columns show only cells located in the MRN, and each column shows a different supertype level cell type in green. (**b.**) Supertype level cell type proportions for top fifteen most abundant types across the MRN subregions. Cell types visualized in (**a**.) are denoted with green asterisks. (**c.**) Selected 4 differentially expressed genes across regions. Each gene is expressed at least log fold change greater than 1 relative to the other domains. MERFISH probe distributions for select genes indicated with pink asterisks are shown in (**d.**). (**d.**) Gene expression gradients across tissue section 25 for *Pax5*, *Six3*, and *Bnc2*, showing specificity for each of the putative MRN subregions. Intensity of color is 0-1 normalized after log scaling raw probe counts. Each dot is a cell, and the color shows the relative transcript count. We show only cells within the subregions to make it visually easier to distinguish the relevant cells. (**e.**) Bar plot of the percentage of cells for a given neurotransmitter type found in each domain, (GABA - GABAergic; Glut - Glutamatergic; NN - non-neuronal). (**f.**) Number of unique cell types (at supertype level) found in each domain, grouped by neurotransmitter class..

We again visualized the neurotransmitter composition and number of unique cell types of given neurotransmitter classes. We observed that the number of types of excitatory neurons was spatially graded. Dorsal domains of MRN (domain 365 shown in purple and domain 198 shown in brown, (**Figure 5e**) featured the highest proportion of glutamatergic neurons, and the proportion of glutamatergic neurons in MRN domains decreased with increasing depth. Nonneuronal cells are also organized along this gradient, but in the opposite direction, the ventral areas featuring the highest proportion of glia and the dorsal the lowest. Interestingly, MRN domains composed of a higher proportion of glutamatergic neurons were also the ones with the greatest number of glutamatergic neuron types, also following a dorsal-ventral gradient (Pearson correlation *r =* 0.89). This relationship was observed for the nonneuronal cells (Pearson correlation *r* = 0.81, **Figure 5f**), but not for GABA-ergic neurons (Pearson correlation *r =* -0.64). This suggests that CellTransformer can identify plausible structures even in historically difficult to characterize areas.

### CellTransformer enables scaling up to multi-animal, million cell datasets and generalizes to other spatial assays

In order to investigate CellTransformer’s ability to integrate across animals, we trained a new model from scratch on the Zhang et al. (2023)^11^ MERFISH data, which uses an 1129 gene panel and is split over four animals, with both coronal (Zhuang 1 and 2) and sagittal sections (Zhuang 3 and 4). We computed embeddings for each neighborhood as in the previous analysis and performed *k*-means, concatenating representations for all mice and sections. This provided an opportunity to examine whether CellTransformer could adapt to a multi-animal case in addition to finding spatial domains across tissue sections of the same animal.

Spatial domains in sequential tissue sections appeared highly concordant across all four mice (**Figure 6a**) at the 50-domain resolution. We used 50 domains to facilitate clear visualization of the domains across animals with relatively few colors. Coronal and sagittal sections across mice clearly corresponded anatomically. Cortical layers were highly consistent across animal and section orientation. Structures that appear in the coronal view can be readily identified in the sagittal sections. For example the hippocampal formation (blue) is well delineated in sections 088 for Zhuang 1, section 044 for Zhuang 2, and across displayed sections of Zhuang 3 and Zhuang 4. Despite a relatively low number of cells in mouse 4 (162,579 cells versus more than 1.5 million for each of the other animals), nearly all spatial domains observed for Zhuang 4 are present in other animals. Note that sections from this animal only cover a section of the lateral portion of the brain and do not span the entirety of the sagittal plane.

**Figure 6.**
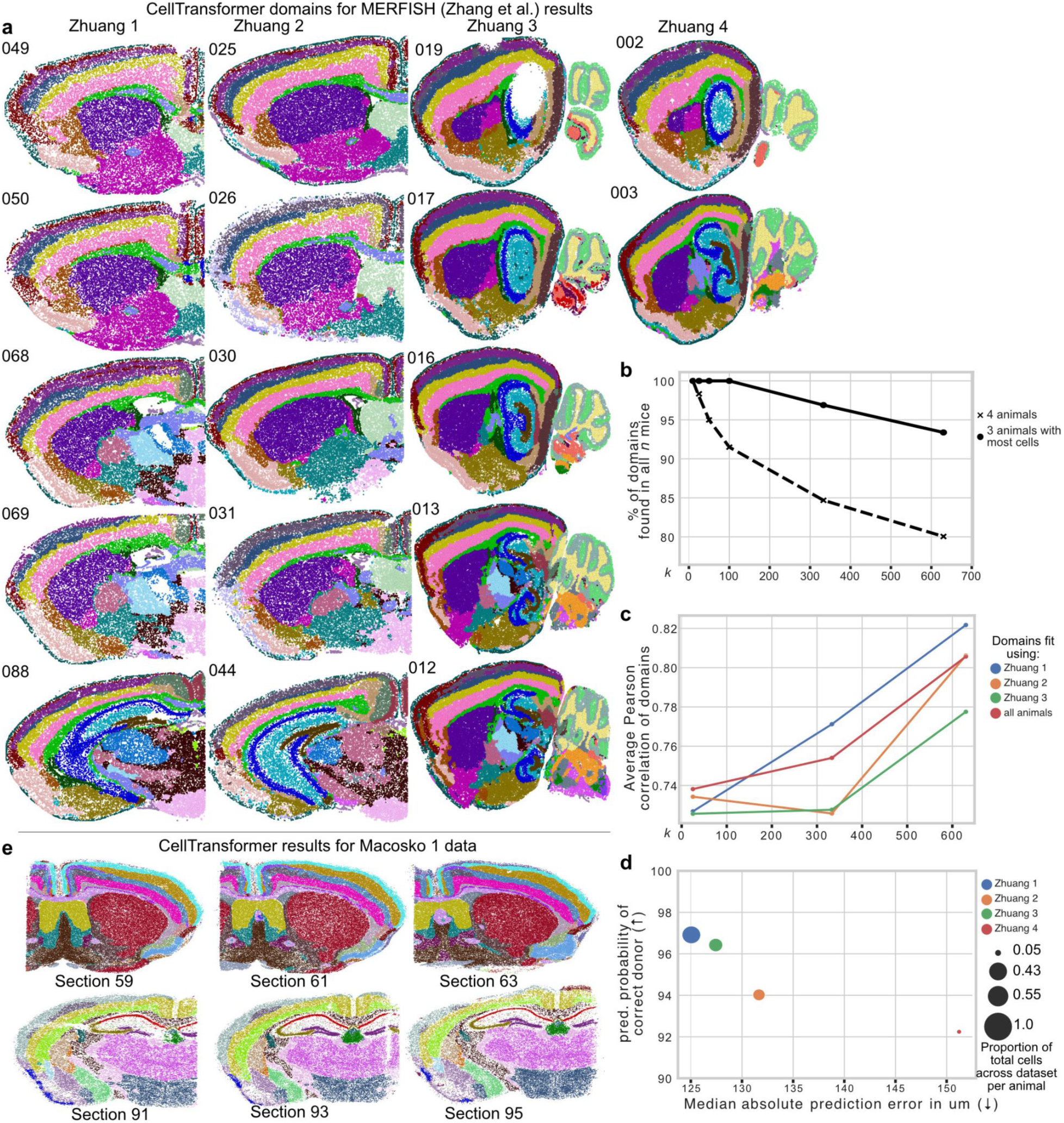
Investigation into performance of CellTransformer on the Zhuang 1-4 datasets (239 sections, both coronal and sagittal, with a 1129 gene MERFISH panel^11^). (**a.**) Representative images of all four mice arranged by column. The section number for each mouse is shown in the upper left of each image. Note that Zhuang 4 only had three sections. For each image, each dot is a cell neighborhood and colors come from a spatial clustering with *k=50* (number of CCF regions at structure level), fit by concatenating embeddings across mice. (**b.**) Quantification of number of per-mouse specific spatial clusters, computed by clustering at different *k* and computing the number of clusters found for all mice (4 animals) and for the three mice with the most cells per mouse (Zhuang 1, 2, and 3). Note that because serial sections were collected at a higher frequency (100µm versus 200µm), different areas of the brain will have marginally higher coverage in one brain or another. (**c.**) Average correlation of the cell type composition of brain regions computed CellTransformer to CCF regions, computed using the linear-sum assignment matching algorithm (exclusively matching regions from one set to the other). Dotted lines with “o” marker indicate results when fitting using all three mice with >1M cells together. Solid lines with “x” marker indicate results when computing spatial clustering on each mouse in isolation. (**d.**) Quantification of subject-level information present in embeddings using linear regression. The median absolute prediction error (x-axis) quantifies accuracy in predicting the (x, y, z) coordinates of a neighborhood from its embeddings. The y-axis quantifies accuracy when predicting mouse identity from embeddings using logistic regression. Values are averaged across cells per mouse. (**e.**) Results of domain discovery (*k*=50) on a Slide-SeqV2 whole mouse brain (Macosko 1) dataset^12^. Two sets of three sequential sections are shown.

We quantified the robustness of CellTransformer domains in a multi-animal context across and within Zhuang 1-4 datasets. We ran clustering and identified domains at the three values of *k:* 25, 333, 630. These *k* values correspond to three CCF resolution levels reported by registration in Zhang et al. (2023) (note the number of domains differs due to registration differences). For each *k* value, we counted the number of domains observed in all four animals. We also repeated this analysis without data for Zhuang 4, which contains far fewer cells than the datasets from other animals (**Figure 6b**). We find that even at high resolution (630 domains), 93.3% domains were found in each mouse, showing high consistency of CellTransformer domains across datasets. With the Zhuang 4 included, at 630 domains, 80.0% domains were found in every animal. To verify that domain consistency across animals was not related to loss of domain spatial coherence, we repeated the neighborhood smoothness analysis we developed for analysis of the Allen 1 dataset on the combined Zhang et al. (2023) data. Spatial smoothness was similar to that of Allen 1 (**Supplementary Figure 10**), indicating CellTransformer can discover spatially coherent domains that are robustly integrated across animals.

We next quantified the similarity of CellTransformer domains to CCF regions. Similarly to our analysis of the Yao et al. (2023) dataset, we computed average similarity of cell type composition vectors from CCF and CellTransformer. In domain discovery across all animals, we found cell-type composition vectors that correspond strongly to CCF (Pearson correlation = 0.805, red line in **Figure 6b).** We also evaluated whether clustering only on embeddings from one animal would significantly affect similarity to CCF. Correspondence between CellTransformer domains and CCF is high even when domains are fit with a subset of the dataset (Pearson correlations > 0.7 for all comparisons across resolutions and domain source, **Figure 6c**). This demonstrates CellTransformer can reproduce a consistent neuroanatomical structure even with a small number of observations. Results were highly similar overall to CCF in the Zhang dataset and Yao dataset (Pearson correlations greater than 0.6 for all comparisons), indicating CellTransformer’s robustness to changes in gene panel and preprocessing choices.

To further investigate how donor metadata was encoded in the embeddings, we employed linear probing strategies commonly used in interpretation of deep learning model embeddings. We regressed CCF-registered (x, y, z) coordinate position across all embeddings and used logistic regression to classify animal identities. Neighborhood level prediction of donor identity was very accurate (>94% for all animals, **Figure 6d**) and median absolute prediction error was accurate within 151 µm. Decoding accuracy for both metrics was strongly associated with the number of cells (Pearson correlation -0.92 for coordinate error and 0.92 for predicted donor probability). The observation that mouse donor identity is easily predicted from per-neighborhood embeddings while still maintaining cross-animal and cross-section coherence is another demonstration of the richness of the representation learned by our approach.

Finally to demonstrate the applicability of our strategy to a different spatial transcriptomics modality, we analyzed a whole mouse brain Slide-SeqV2 dataset (“Macosko 1”), collected in Langlieb et al. (2023)^12^. Slide-SeqV2 provides whole transcriptome coverage in a spatial context by tiling tissue slices with 10 µm by 10 µm squares. As each square may contain more than one cell or a partial cell, we fit our model to the deconvoluted single-cell data computed using the RCTD^32^ method provided. This produced 4,783,976 cells across 101 slices. We also filtered the dataset for low-quality cells and infrequently expressed genes (see **Methods**). We found that increasing the size of our model (from 4 encoder layers to 10) was necessary to identify spatially coherent domains, perhaps driven by the much larger number of genes detected (5019 versus 500 or 1129 in the two MERFISH datasets; see **Methods**). We plot three sequential sections from domain discovery at *k*=50 (**Figure 6e**). We show that CellTransformer robustly identifies cortical layers across sections and in known structures such as the midbrain and the piriform areas. Domain discovery with greater than 50 regions did not produce adequate integration across sections, possibly because of variable cellular density and single-cell read depth across sections.

Overall, we found that the CellTransformer workflow successfully identifies interpretable domains across different spatial transcriptomics modalities, and that the resolution of cross-section and cross-dataset domains depends on the specific spatial transcriptomics method and the quality of the datasets.

## Discussion

In this study, we present a transformer-based pipeline to combine scRNA-seq and spatially resolved transcriptomic atlases to perform accurate organ level domain discovery. We employed a novel representation learning workflow and implemented a computationally efficient pipeline that readily allows scaling to multi-million cell, multi-animal datasets. The representations learned in our model can be clustered to identify progressively finer-scale spatial domains directly from local cellular and molecular information alone, without predefined spatial labels. We show these regions can be interpreted at the gene or cell level and recapitulate a variety of existing findings in the neuroscience literature where many existing methods cannot. Our pipeline allows extraction of a very high number of domains, which retain high correspondence to existing brain region ontology (CCF). These domains are also highly spatially consistent both within and across tissue sections and even over multiple animals. To our knowledge this is the first demonstration of fine-grained domain detection beyond human annotation (CCF) using data-driven methods in transcriptomic data. Not only can CellTransformer discover this fine structure, but it can reliably find it across animals even with hundreds of regions. This capability is intrinsic to our model, and is learned despite any conditional modeling for donor or section-level covariates, indicating the robustness of learned features.

We demonstrated the robustness of our model in uncovering biologically relevant models and characterized our pipeline’s ability to reproduce known neuroanatomy in the hippocampal formation and superior colliculus. Detected domains were concordant with previous comprehensive transcriptomic and connectivity studies of these areas but were identified in a scalable and data-driven way. Not only were we able to detect known and novel regions, we found that CellTransformer domains can recapitulate and extend known spatial cell type enrichment patterns and gene expression gradients.

We highlight several advantages of our architecture and approach. Although the use of graph-structured architecture or self-attention to model cells in a neighborhood graph is not novel^14,21,33^, our approach combines novel self-supervised training objective based on spatial correlation between a cell and its neighbors that facilitates learning of a fixed representation for a cellular neighborhood (**Supplementary Note 2**). The intersection of graph neural networks, transformers, and representation learning research is a rich and rapidly moving research area. Methods for spatial-graph structured data such as CellTransformer will benefit immensely from implementing more effective ways of encoding the data and its metadata such as better position encoding mechanisms^34^, rotationally-invariant architectures^35^, or arbitrary numbers of genes^36^. There are also significant opportunities to extend CellTransformer’s local representation framework to include other data modalities. Using a transformer than graph neural network facilitates inclusion of arbitrary contextual data such as cell-level (e.g. neurophysiology^37,38^) and pixel-level data (e.g. mesoscale axonal connectivity^39^, or magnetic resonance imaging^40^) which can be tokenized and included in our framework.

Caveats of our approach include the necessity of a user-specified spatial radius (for neighborhood computation) and choice of *k* for spatial cluster detection. The stability of detected spatial domains at a given radius or *k* poses an interesting future angle from which to study anatomical hierarchies in the brain. Users must also have access to GPUs (to allow for timely model fitting), which reduces overall accessibility, although the hardware requirements are still much less intensive than for many existing models such as spaGCN and scENVI^4^.

CellTransformer advances the state of the art for automated domain detection by allowing identification of granular and biologically relevant spatial domains that is extensible to both very large, multi-animal spatial transcriptomic datasets. As spatially resolved transcriptomic and multi-omic studies of the brain become more prevalent, tools such as CellTransformer provide avenues to transform data into refined anatomical maps of the brain and other complex organs and pave the way towards tissue-level structure-function mapping.

## Methods

### Allen Brain Cell Mouse Whole Brain (ABC-MWB) dataset processing

#### Allen Institute for Brain Science dataset preprocessing

We downloaded the log-transformed MERFISH probe counts and metadata for the Allen Institute for Brain Science animal (“Allen 1”) from the Allen Institute public release (https://alleninstitute.github.io/abc_atlas_access/intro.html) access for ABC-MWB.The Allen 1 dataset is composed of 53 coronal sections. The MERFISH probe set included 500 genes. Serial sections were collected at 200 μm intervals. We used the taxonomy from the “20231215” data release. Allen 1 is composed of 3,737,550 cells. We transformed the (x, y) coordinates of each cell into microns instead of mm as provided. Otherwise the dataset was used as-is for neural network training.

#### Zhuang lab (Zhang et al.) dataset processing

Data were downloaded from the “20230830” data release from the Allen Institute ABC-MWB public data release. Two animals (“Zhuang 1” and “Zhuang 2”) were collected with coronal sections. The other two animals (“Zhuang 3” and “Zhuang 4”) were collected sagittally. Serial sections for Zhuang 1 (female) were collected at 100 μm intervals, while serial sections for the other animals (all male) were collected at 200 μm intervals. The size of the MERFISH probe set included 1129 genes. Zhuang 1 and Zhuang 2 consist of 2,846,909 cells and 1,227,409 cells, respectively. Zhuang 3 and Zhuang 4 consist of 1,585,844 cells and 162,579 cells, respectively. We transformed the (x, y) coordinates of each cell into microns instead of mm as provided. Otherwise, the data were used as-is for neural network training.

#### Cellular neighborhood construction

We consider cells in the same neighborhood as a reference cell if the distance between them is within a box of fixed size. For all MERFISH datasets we used a box width of 85 μm.

#### CellTransformer architecture

We construct a CellTransformer to generate a latent representation from a cellular neighborhood where this representation is composed of both molecular and cell type information. We represent cells as nodes in an undirected graph, 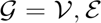 where *V* indexes the nodes in the graph (cells) and we add an edge 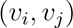 to the edge set ε if *d_i, j_* < *r*, with *r* a user-specified distance in microns. We assume also that for each node we have access to 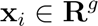, a *g*-dimensional vector of MERFISH probe or cell deconvoluted transcript counts. We also assume we are given class labels 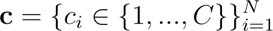 for each of the *N* cells. The user must also specify an embedding dimension and number of transformer encoder and decoder layers; in all experiments in this paper using MERFISH data we use an embedding dimension of 384, 4 encoder layers, and 4 decoder layers. For Slide-SeqV2 analysis we used 10 encoder layers and 4 decoder layers.

To generate a neighborhood embedding, we identify a particular cell which we call a reference cell. Its first degree neighbors are extracted from *G*. We first apply a shallow encoder (two layer perceptron with GELU nonlinearity) function 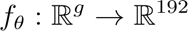 which maps the gene expression into embedding space. We likewise construct and apply the function 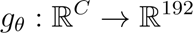 to map one-hot encoded cell type labels to embedding space, here a simple lookup into a learnable embedding table. These representations are concatenated into a single 384-dimensional representation. We apply these transformations to each cell in the neighborhood, not including the reference cell. We note that at this point, all operations have been per-cell without interactions. In addition to these cell tokens, we also instantiate for each neighborhood a register token which we use to accumulate global information across the neighborhood. We refer to this token as a <CLS>-like token in keeping with previous literature.

We then apply a transformer encoder to the cells, only allowing cells within the same neighborhood and their <CLS>-like tokens to attend to each other. We use 8 attention heads with GELU activations and layer norm prior to attention and MLP projection. We note that including a bias term in the key, query, and value MLPs is important to stabilize training, while not noting any significant differences in models fit with and without bias terms for the rest of the encoder and decoder layer MLPs. Following the transformer encoder, we use attention pooling to aggregate the cell and <CLS> representations for each neighborhood into a single token with embedding dimension 384. We refer to these as the neighborhood representations.

We then instantiate a new token from each reference cell that is a learned embedding for each cell type (separately from the encoder cell type embedding). These are concatenated to the neighborhood representations. We then apply a transformer decoder to the tokens, allowing only the neighborhood token and masked cell embedding to attend to each other if they are from the same cellular neighborhood. This decoder embedding dimension was 384 with 8 attention heads.

During training, we extract only the masked reference cell tokens. We then use separate linear projections to output the mean, dispersion, scale, and zero inflation logit parameters for zero-inflated negative binomial regression. We optimize the model by minimizing the log likelihood of a negative binomial distribution using observed cells’ MERFISH probe counts. We trained all versions of CellTransformer on a system with 2 NVIDIA A6000 GPUs with effective batch size 256.

#### Spatial domain detection

Once trained, we apply CellTransformer to a given dataset and instead of extracting reference cell tokens we extract the neighborhood representation. We then cluster this representation using *k-*means. We use the cuml library to perform this operation on GPU (cuml.KMeans), with arguments n_init=3, oversampling_factor=3, and max_iter=1000.

#### Optional smoothing of embeddings

We observe spatial domains are spatially smooth. However in the case that there is a high-frequency signal that the end-user would like to filter, we optionally introduce a step prior to *k-*means where we smooth the embeddings using a Gaussian filter. For all comparisons except those in **Supplementary Figure 12**, smoothing was performed with a Gaussian filter with 40 micron full-width at half maxima (sigma of 12.01 microns).

#### Model fitting on the Allen 1 dataset

We used an 80%-20% train-test split proportion (random splitting across the entire dataset) and the ADAM optimizer over 40 epochs. We perform a linear warmup for 500 steps to a peak learning rate of 0.001 and use an inverse-square root learning rate scheduler to decay the learning rate continuously. We use a weight decay value of 0.00005 which we do not warm up.

#### Model fitting on Zhuang datasets

We perform training from scratch without transfer. We trained for 40 epochs with the same settings as for the Allen 1 with the exception of adapting projections to 1129 genes instead of 500.

#### Computation of stability criterion

We follow Wu et al. (2016) in using an Amari-type distance to compare clustering solutions. Briefly, we compute several replicates (20 in this work) of *k-*means at a given choice of *k* with different random seed as, **D***_k, i_* with *i* indexing the different centroids for a given solution. We then measure the stability of a given choice of *k* by comparing the similarity of all pairs of *D_k_*. Define C the Pearson correlation matrix between pairs D and D’. Then we use this dissimilarity metric:

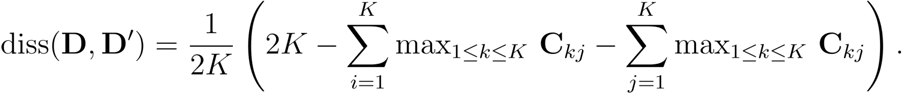

to identify the choice of *k* which is most stable.

#### Regional matching with CCF computation

To quantify overall similarity of regions extracted using CellTransformer with CCF, we first extract cell type composition vectors for each region at a given level of the hierarchy. For all comparisons in **Figure 2**, we use the subclass level (338 cell types), resulting in *k-*region by 338 matrices. For each region derived from one of the tested models, we compute two quantities: the best match (maximum value of Pearson correlation, non-exclusively) to any CCF (**Figure 2d**) or an exclusive match (using the linear sum assignment algorithm) to pair the regions from either set one-to-one (**Figure 2e)**. We then computed the average Pearson correlation across the paired matches as the metric. We use scipy.optimize’s implementation to solve the linear sum problem.

#### CellCharter

To run CellCharter we first generated scVI embeddings using the default settings for depth and width of the network and with the tissue section labels as conditional batch variables. We trained for 50 epochs using the early_stopping=True setting. We then aggregated across 3 (default settings), 6, 9 layers using the cellcharter.gr.aggregate_neighbors function. We then applied CellCharter’s Gaussian mixture model implementation at various choices of the number of Gaussians. We could not run the mixture model with our hardware (A6000, 48GB GPU memory) for more than 9 layers, which was also the number which produced the highest correspondence with CCF and is reported in **Figure 2e**.

#### SPIRAL

To run SPIRAL we generated edge sets for 40um, 85um, and 170um neighborhood radii. SPIRAL requires supervision on single-cell types so for this we use the subclass cell type levels. We trained models across neighborhood sizes for 1 epoch and then chose the neighborhood size with best performance (170um) and trained this model to saturation (10 epochs). SPIRAL uses four objective functions so to assess saturation we averaged them. We note that SPIRAL does not use a training and testing set split in their training, making it difficult to assess an optimal stopping point. For the *k*=354 and *k*=670 domain discovery analyses the SPIRAL clustering pipeline produced an out-of-memory error and we instead used our own pipeline with *k*-means on SPIRAL embeddings.

#### Nearest-neighbor smoothness computation

To quantify smoothness of the spatial domains, we use a nearest-neighbor approach. We extract approximate spatial neighbors for each cell using cuml.NearestNeighbor with 100 neighbors, restricting neighbors to be within the same tissue section. For a given domain set, either from CCF, CellTransformer, or CellCharter, we extract the spatial domain label of the given cell and count the proportion of times that cell is observed in the 100 neighbors. These proportions are averaged across all cells and tissues.

#### Linear probing experiments

We extract neighborhood representations for each of the cells in the Zhuang lab datasets. First, we regress these embeddings on the (x, y, z) coordinates. We then computed the absolute prediction error in terms of the coordinates and then reported the average. We also fit a multi-class logistic regression using the mouse donor identity. For the logistic regression we use cuml.LogisticRegression with default settings in cuml. For the cell position regression we fit simple least squares using PyTorch via QR decomposition.

#### Quantification of spatial contribution to gene expression

We interpret the accuracy of the gene expression predictions for a given cell as an index of correlation of an instance of a particular cell type with its surrounding neighbors. To do this, we compute a simple baseline model which predicts average gene expression (computed across the entire Allen Institute for Brain Science mouse dataset) for each cell. We compute the average Pearson correlation for each instance of a given cell type and average across instances to obtain an average Pearson correlation. We then compute a Pearson correlation between each cell’s observed gene expression and the CellTransformer predictions, averaging similarly across instances of a given cell type. The difference between the baseline and model predictions is displayed, per cell type, and grouped across neurotransmitter types in **Supplementary Figure 12**.

#### Zhuang lab dataset per-animal CCF comparison

We contrast two methods of extracting spatial domains from the four animals in the Zhuang lab dataset^11^. We first fix *k,* the number of desired spatial domains. Then we fit one *k-*means model on all of the neighborhood embeddings for all four (Zhuang 1, 2, 3, and 4) mice together. We also fit a *k-*means model to the embeddings of the mice separately. We then compute the similarity of these region sets using the same method used to quantify differences between CellCharter and CellTransformer by comparing their regional cell type composition vectors.

#### Slide-seqV2 analysis

Initial results with a direct transfer of hyperparameters to the Langlieb et al. (2023) dataset^12^ did not produce spatially coherent domains. We therefore implemented two quality control procedures on the raw data. After filtering for coding genes and non-mitochondrial genes, we additionally used only genes that were expressed in >10% of cells in the dataset. At the cellular level we identified cells with >20% mitochondrial genes and those within the 10th percentile of read depth across each section. We also removed these cells. This left 5,019 genes and 4,783,456 cells. We noted that a successful segmentation in the Langlieb et al. (2023) dataset required a larger model than the MERFISH ones, using 10 encoder layers rather than 4, which we attributed to the 10X higher number of genes in this dataset versus the 500 in the ABC-MWB Allen 1 dataset. We used a neighborhood size of 50um to reduce memory footprint, reasoning the higher cell density in this dataset and higher number of genes would provide enough information for representational richness.

#### Other software

Principal software used in this work includes PyTorch^41^, numpy^42^, scikit-learn^43^, scipy^44^, scanpy^45^, cuml^46^, matplotlib^47^, and seaborn^48^.

## Code and data availability

Code will be publicly available at https://github.com/abbasilab/celltransformer. All data used for this publication is available either at the Allen Institute ABC-MWB data portal (https://portal.brain-map.org/atlases-and-data/bkp/abc-atlas) or the CZI cellxgene portal (https://cellxgene.cziscience.com/datasets).

## Acknowledgments

The authors would like to acknowledge Forrest Collman for help with Neuroglancer data visualization of spatial domains. We acknowledge Patrick Xian for helpful discussions. AJL acknowledges Katharine Z. Yu, Chang Kim, Parker Grosjean, Lee Rao, and Tom Nowakowski for feedback on neuroscientific and machine learning contexts of paper. RA, BT, AJL, and HZ would like to acknowledge support from the Weill Neurohub through the Weill Neurohub’s Next Great Ideas Award. RA would like to acknowledge support from the National Institute of Mental Health of the National Institutes of Health under award number RF1MH128672 and Sandler Program for Breakthrough Biomedical Research, which is partially funded by the Sandler Foundation. AJL acknowledges support from the UCSF Discovery Fellowship and the UCSF Graduate Research Mentorship Fellowship. Sharing of Allen Institute for Brain Science data through Allen Brain Cell Atlas (and related tools) and registration of the AIBS MERFISH brain to the CCFv3 was funded through 1U24MH130918-01 to L.N.

## Supplementary Note 1: Effect of smoothing and analysis on striatal glial populations

When scaling up the number of regions past 500 in the Allen 1 dataset, we observed that almost all spatial clusters were spatially smooth except for a recurring pattern in the striatum. We plot (**Supplementary Figure 11a-b**) six sequential sections where we identified an irregular (which we define here as a broadly non-convex shape that does not form relatively singular connected component) pattern of cells in the striatum and only the striatum (note the spatial uniformity of areas surrounding striatum in cortex and endopiriform area, nucleus accumbens etc.). We identified cells in these areas and found they were mostly non-neuronal, with astrocytic types (such as 1163 **Astro-TE NN_3, Supplementary Figure 11**) forming a large proportion of cells.

We sought to understand whether these spatial clusters might be biologically relevant or somehow related to noise. A recent paper, Ollivier et al. (2024)^49^ identified a novel population of *Crym*+ astrocytes in a similar spatial distribution as observed in our regions, specifically in a dorsoventral and lateromedial distribution (see **Supplementary Figure 11e** for reproduction from Ollivier et al., granted with permission). As *Crym* was included in the MERFISH panel, we quantified *Crym* expression in astrocytes within these areas, finding that all but two of these spatially irregular domains had very high levels of *Crym* expression. Notably, the two groups with lower expression, spatial clusters 457 and 758, were the most dorsolateral, and are distributed where *Crym+* astrocytes were not observed in Ollivier et al. We reasoned that these spatial clustersmay have biological relevance.

However, to simplify downstream analyses and conform with neuroanatomical conventions we applied a simple smoothing operation (see **Methods**), which removed this spatial cluster in successive clustering operations. We used a very small smoothing window (12 micron sigma, or 40 micron full-width at half-maxima) and found the order of ranked methods and their relative performance changes were not significantly affected.

## Supplementary Note 2: Interpreting the CellTransformer objective as a measure of spatial dependence

One interpretation of the CellTransformer architecture is learning two representations of cellular gene expression. The first (the learned embedding for each cell type) is unconditional on the spatial neighborhood information. The second is one that is conditional on the spatial neighborhood information learned in the encoder portion of the network and parameterized as a residual update. This residual update can then conveniently be aggregated across layers and represented as a single update term on the unconditional representation to produce the final output. We interpret the increase in accuracy from neighborhood-conditional gene expression prediction as an index of spatial dependence. A trivial or poorly fit optimization solution would produce a small value of this index. A similar idea has been previously presented in a number of works, most recently the NCEM approach^14^.

When analyzing the Allen 1 dataset we observe increases in the predictive accuracy (mean 0.10 +/-0.0701 in correlation, averaged across cell types) across the dataset. Moreover, there are few cell types for which there is a decrease in predictive accuracy, indicating that our model has nontrivially learned the objective (**Supplementary Figure 12a**). Those that are poorly predicted are often only present in the dataset at very low abundances. Conditioned on cell types with more than 10^2^-10^3^ cells, accuracy has only a mild correlation with cellular density (**Supplementary Figure 12b**) or on number of cells of given type in the dataset. Immature neurons (IMN) are the class which benefits the most from conditional prediction, suggestive of their complex migratory dynamics^1^. Note that we compute accuracies at the subclass level (338 types). Increase in accuracy does correlate strongly with log-number of observations per cell type, (Pearson correlation of 0.71, **Supplementary Figure 12c**), possibly indicating inefficiencies in pretraining.

## Supplementary Figures

**Supplementary Figure 1.**
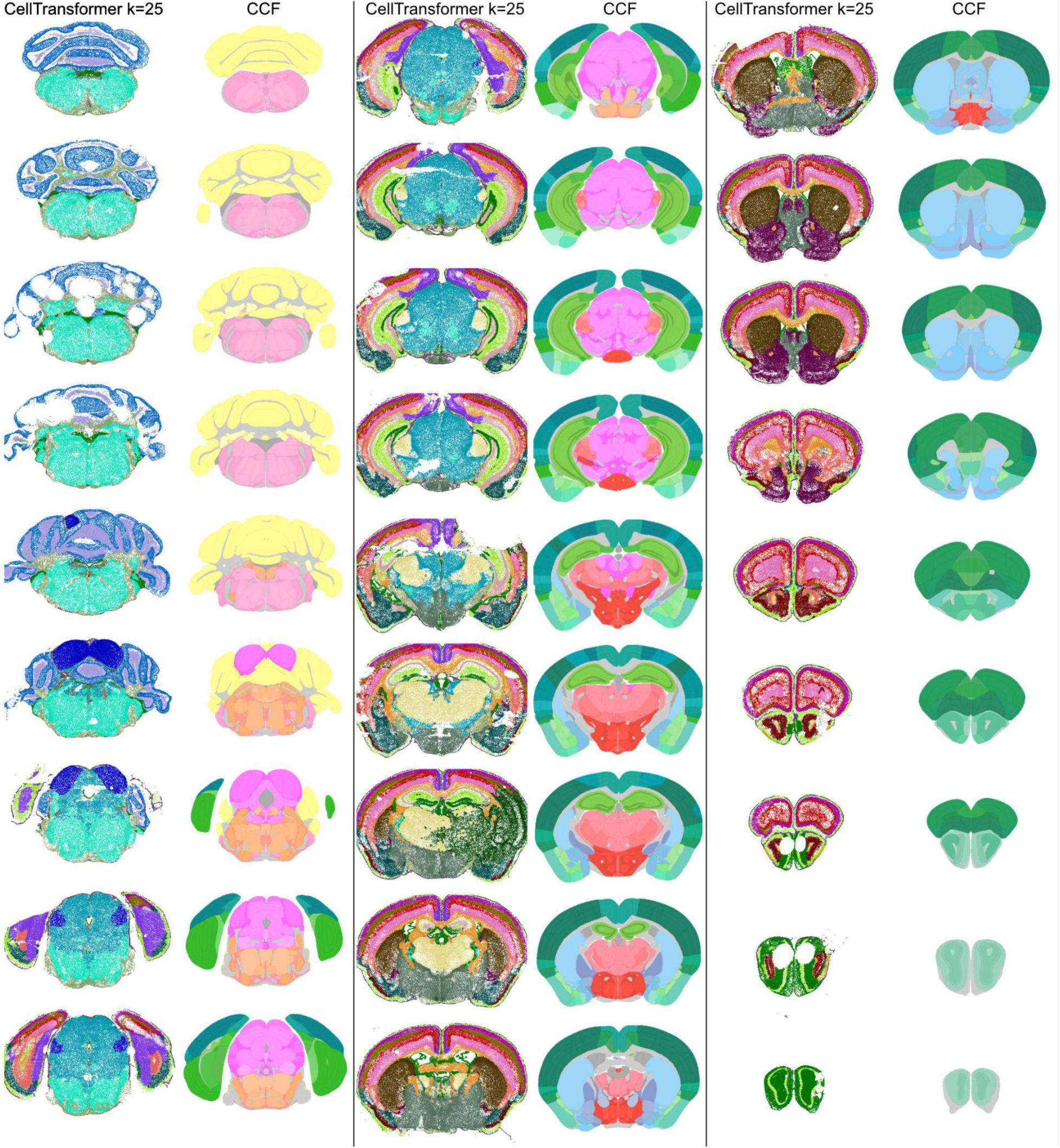
CellTransformer spatial domains (left) and the corresponding CCF annotations (right) organized in 3 columns for roughly half of the sections in the Allen 1 dataset^1^, approximately every other section. CellTransformer domains were calculated at *k*=25 clusters.

**Supplementary Figure 2.**
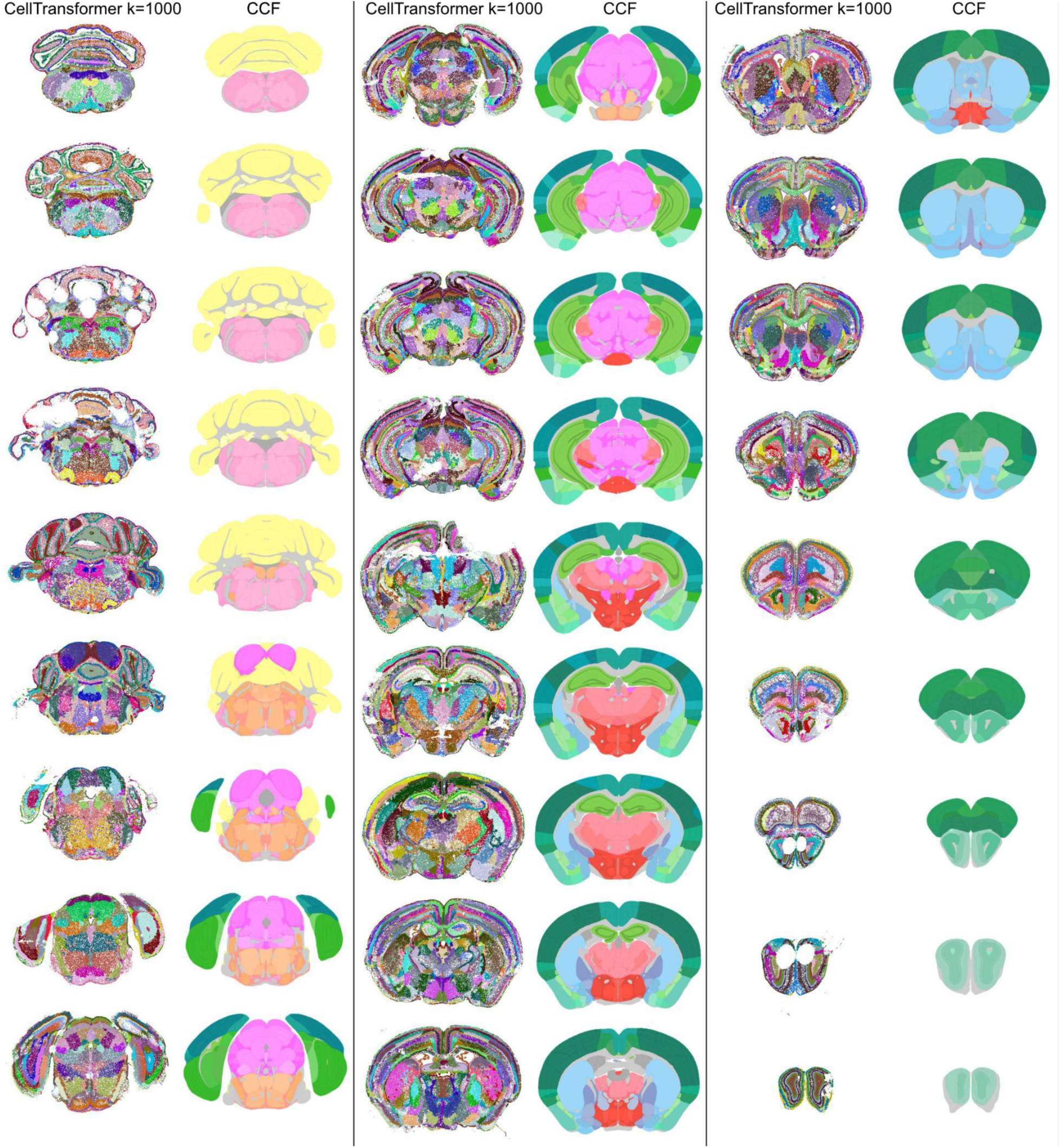
CellTransformer spatial domains (left) and the corresponding CCF annotations (right) organized in 3 columns for roughly half of the sections in the Allen 1 dataset^1^, approximately every other section. CellTransformer domains were calculated at *k*=1000 clusters.

**Supplementary Figure 3.**
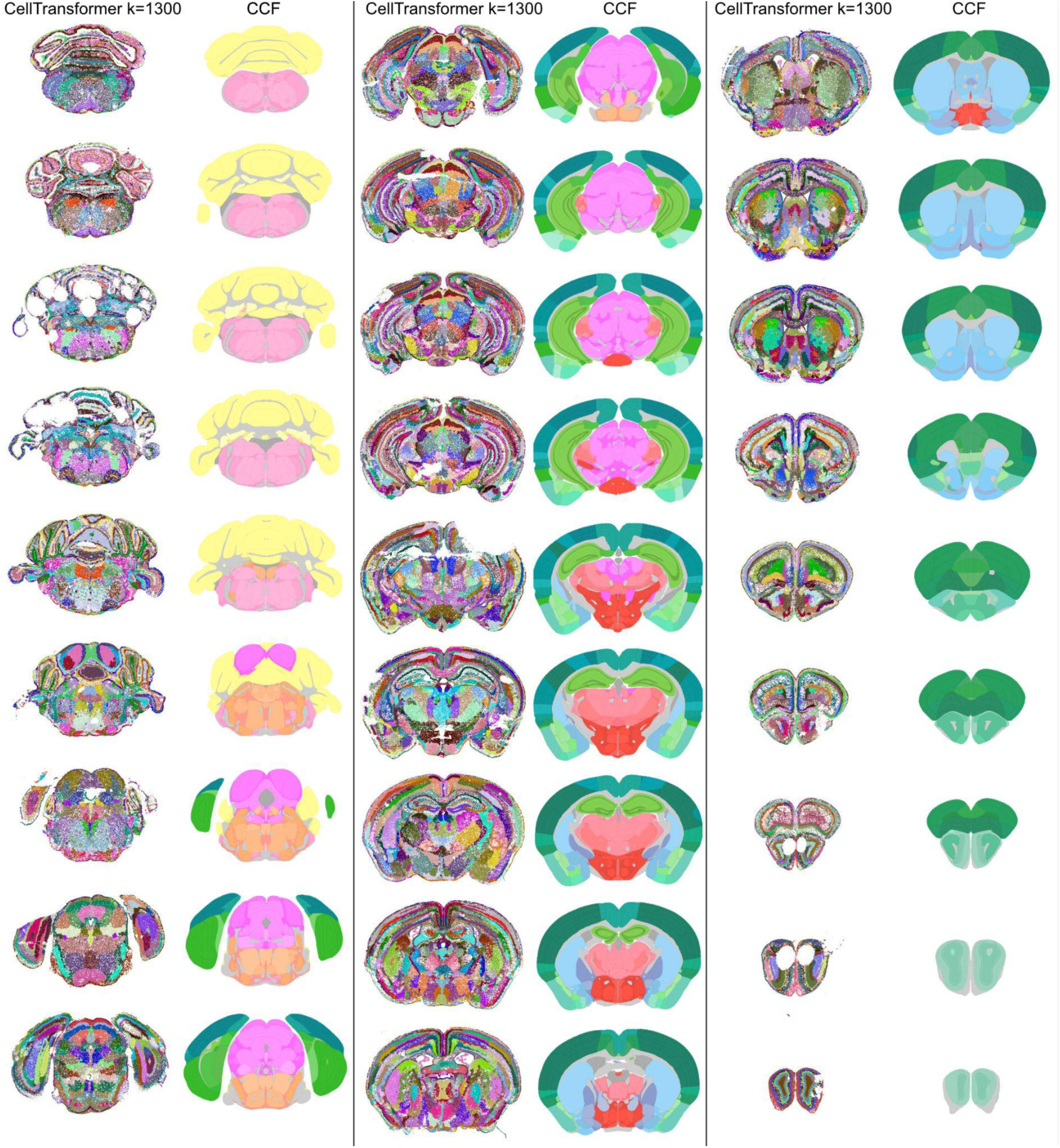
CellTransformer spatial domains (left) and the corresponding CCF annotations (right) organized in 3 columns for roughly half of the sections in the Allen 1 dataset^1^, approximately every other section. CellTransformer domains were calculated at *k*=1300 clusters.

**Supplementary Figure 4.**
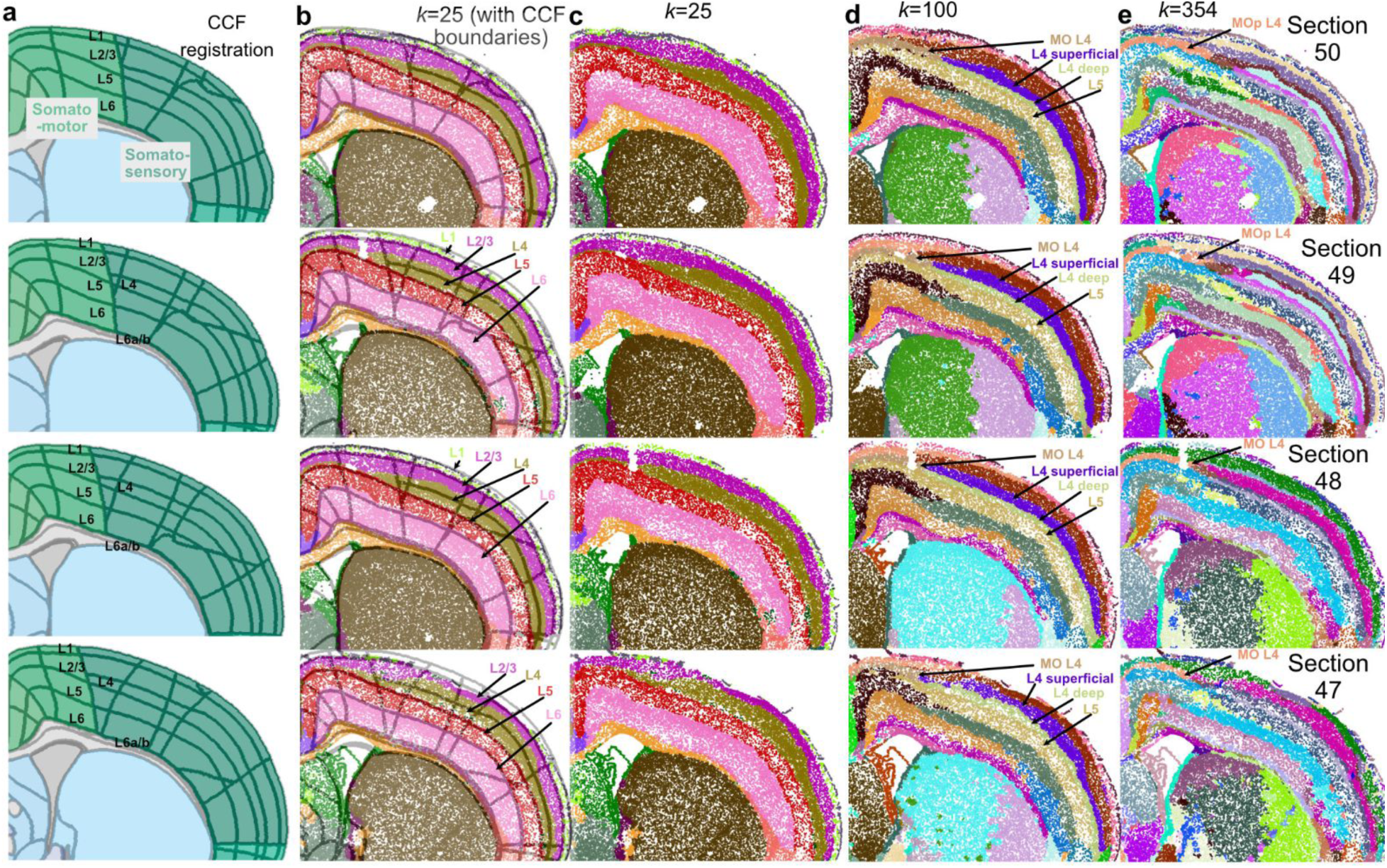
Four sequential sections of the Allen 1 dataset (200 µm sampling interval between sections^1^) displayed with CellTransformer labels at varying resolution alongside CCF registration results. MO: motor cortex. (**a.**) CCF registration of four sequential sections shown in Figure 2. cortical layers are marked based on CCF annotations. (**b.**) *k*=25 spatial domains with CellTransformer shown with regional boundaries from CCF in light gray. Putative cortical layers are annotated, showing CellTransformer replicates known cortical layers. (**c.**) 25 domains shown without CCF annotations to facilitate visualization. (**d.**) Same sections now shown with 100 domains to help show the transition from coarse (25 domains) to fine (100 domains). Sublayers of cortex are identified including layer 4 in motor cortex which transcriptomic studies have verified but has been difficult to identify using histological approaches. (**e.**) 354 domain zoom in on the same sections, showing consistency of layer 4 motor cortex detection as well as an anterior-posterior subdivision across motor and somatosensory cortical layers and clear distinction of cortical layers that lie within motor and somatosensory areas.

**Supplementary Figure 5.**
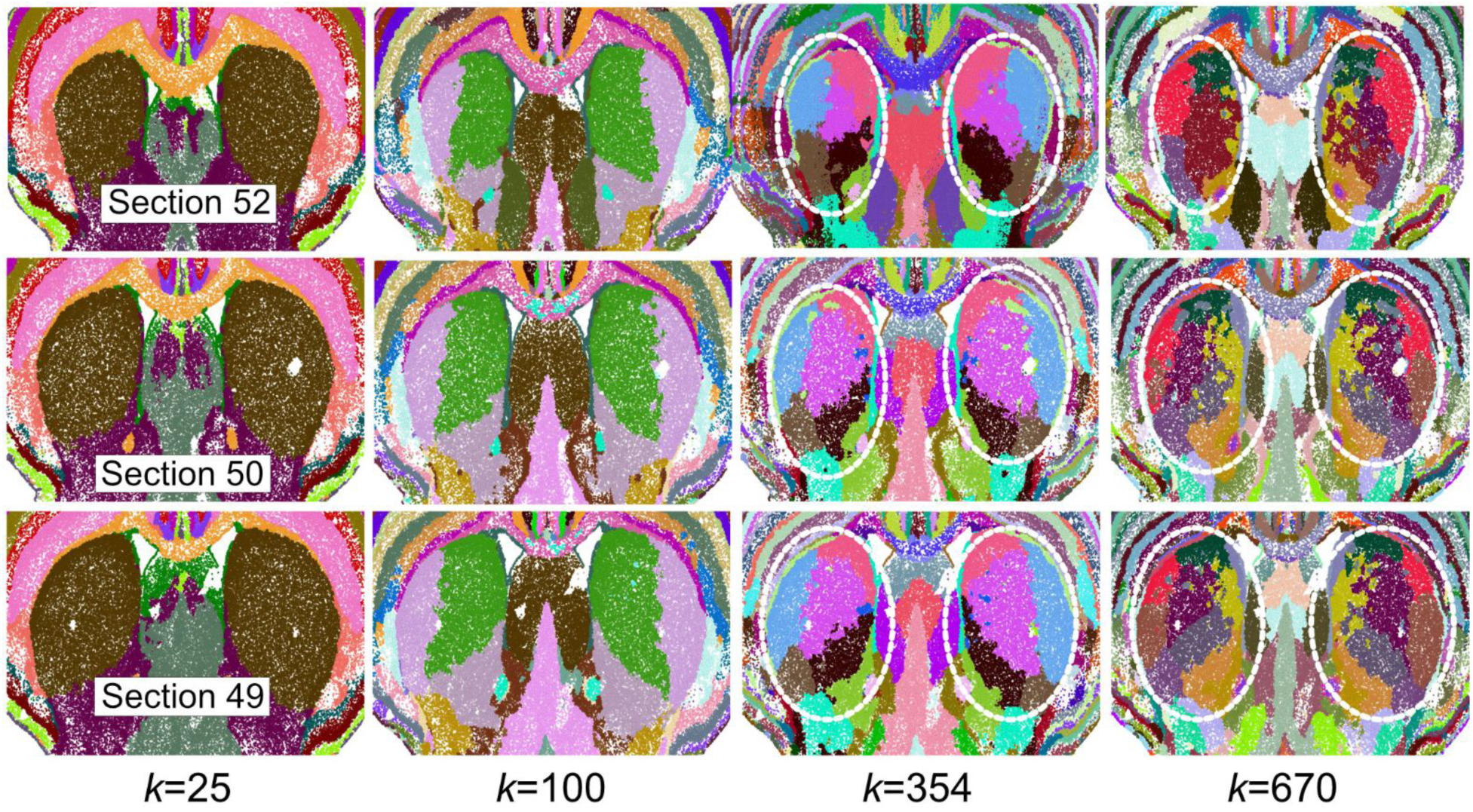
CellTransformer domains identified in the Allen 1 dataset^1^ at varied values of *k*, colored in three sequential sections (200 µm sampling interval between sections with consecutive numbers, top to bottom corresponds to rostral to caudal). Caudoputamen is roughly highlighted by dotted circles in *k*=354 and *k*=670 to assist visualization.

**Supplementary Figure 6.**
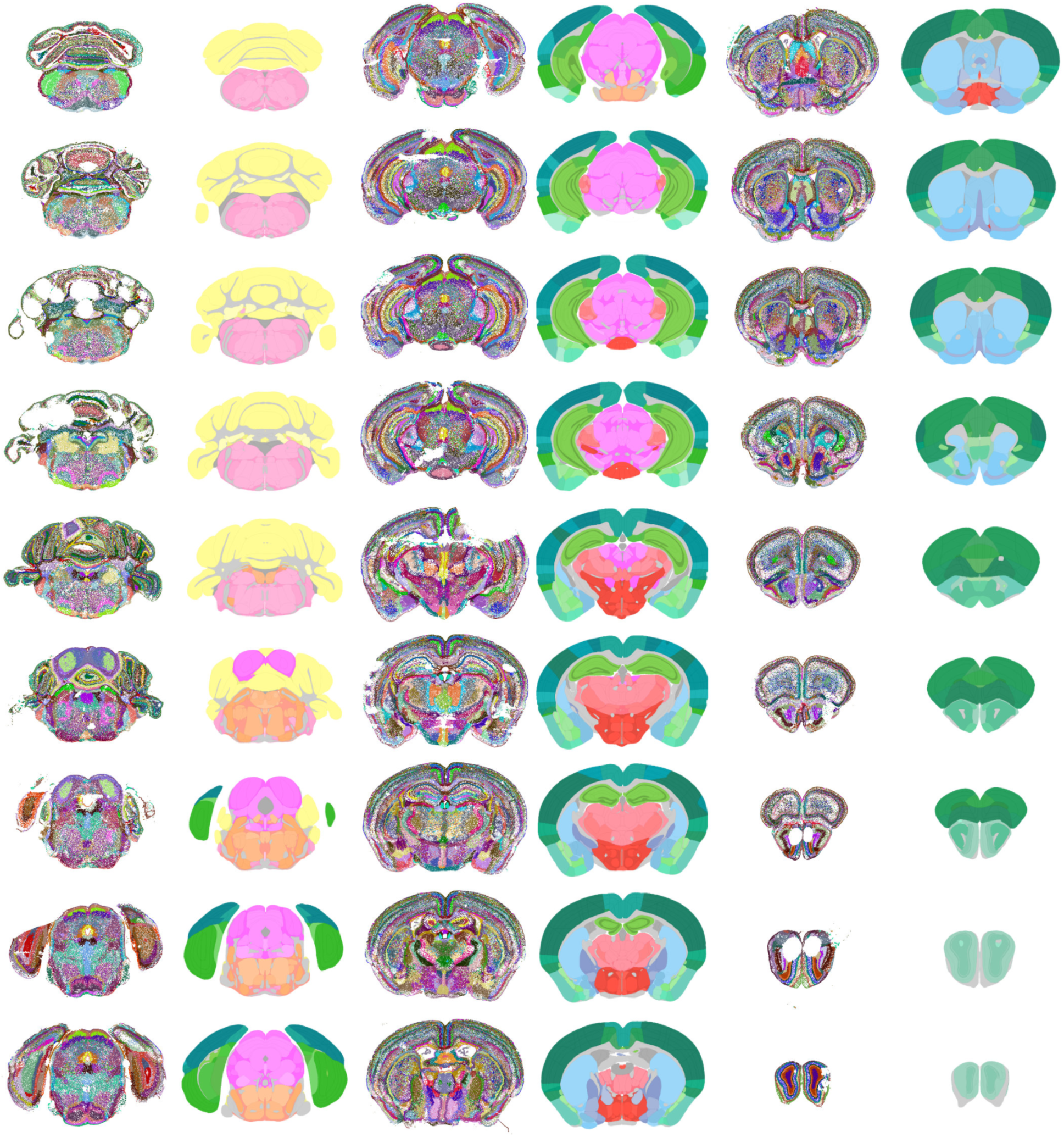
CellCharter results (left) and the corresponding CCF annotations (right) organized in 3 columns for roughly half of the sections in the ABC-MWB (Allen 1^1^) dataset, approximately every other section. The color labels for CellCharter correspond to its Gaussian mixture model implementation with *k=*670 clusters.

**Supplementary Figure 7.**
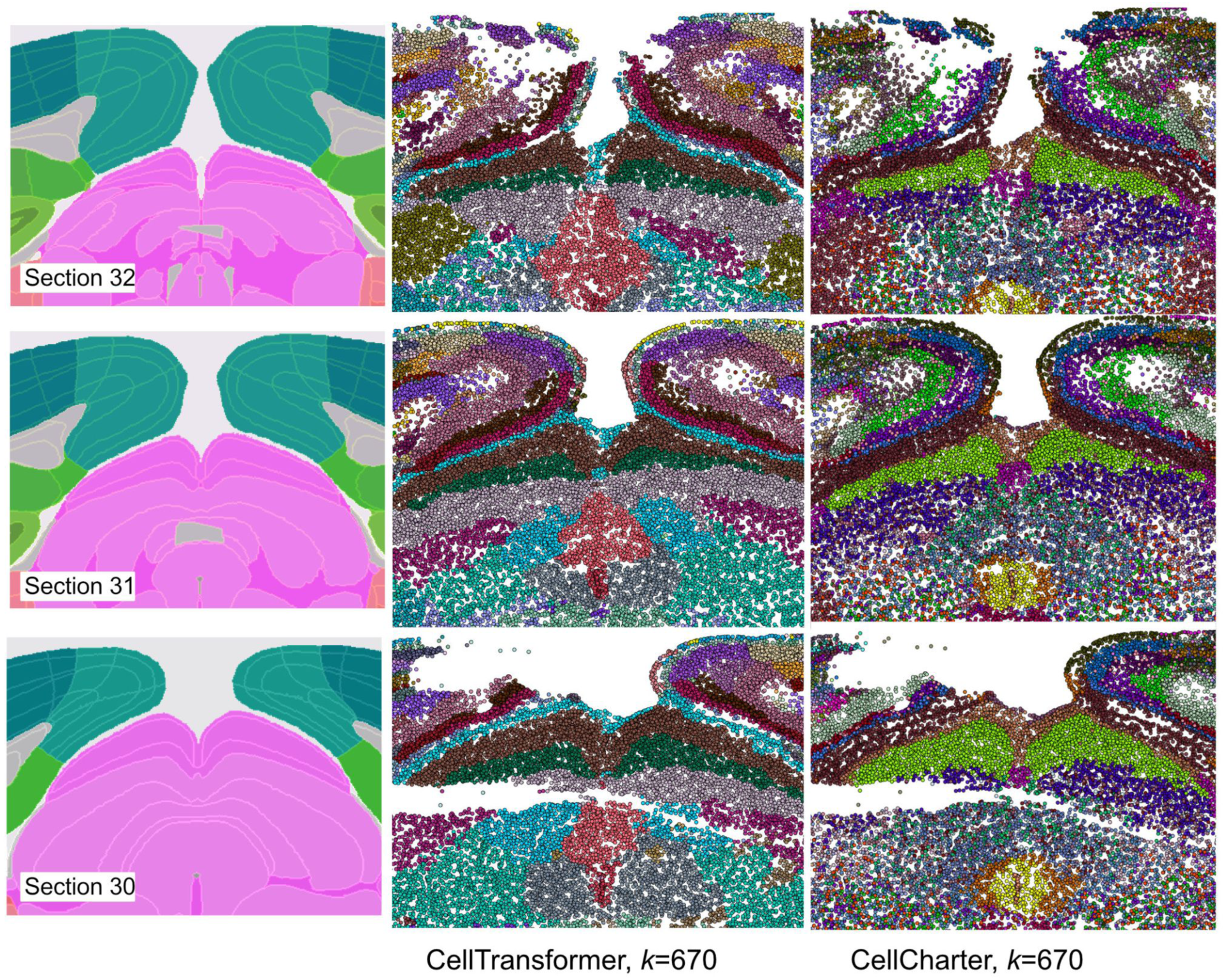
Comparison of spatial domains in midbrain for CellTransformer and CellCharter discovered in the Allen 1 dataset^1^. Left column shows approximate CCF registration. Middle column shows CellTransformer domains at *k*=670 and the right column shows CellCharter domains with 670 Gaussians. The general performance in outlining cortical layers is similar, however in the midbrain, even at half the number of clusters, CellCharter loses spatial coherence compared with CellTransformer. For example, CellCharter only identifies two layers of superior colliculus, whereas multiple layers are defined by CellTransformer.

**Supplementary Figure 8.**
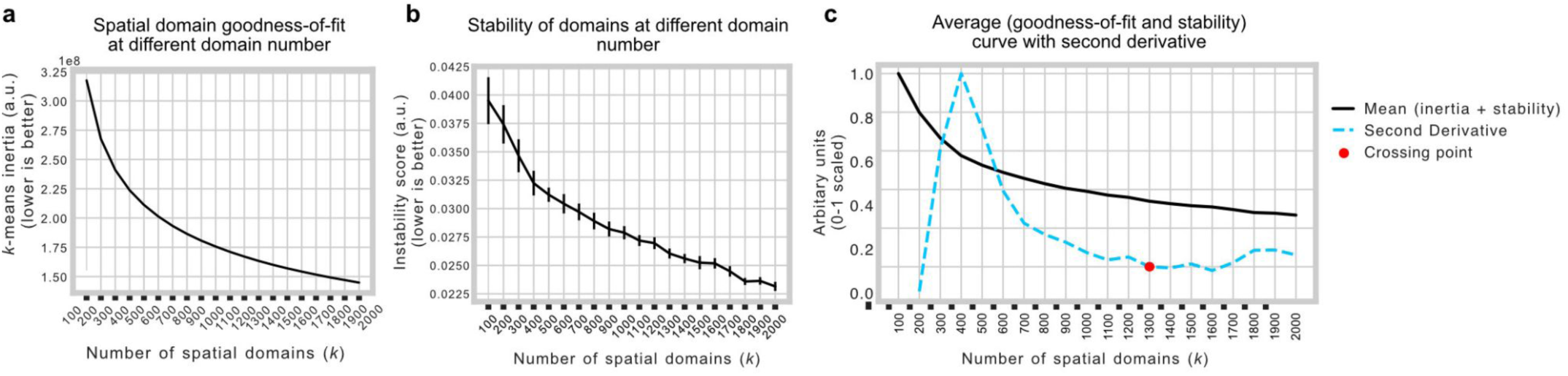
Quantification of goodness-of-fit and stability of varying numbers of spatial domains. (**a.**) inertia (sum of squares errors for each cluster centroid) calculated for different clustering solutions when clustering embeddings generated using CellTransformer on the Allen 1 dataset^1^. Error bars (standard deviation) are calculated but not visible due to scale. (**b.**) instability scores (see **Methods**) calculated for different clustering solutions using the Allen 1 dataset. Error bars are standard deviation. (**c.**) Average of inertia and stability curves (black line) and second derivative of same curve (blue dotted lines). Second derivative crossing point at *k=*1300 shown with red dot.

**Supplementary Figure 9.**
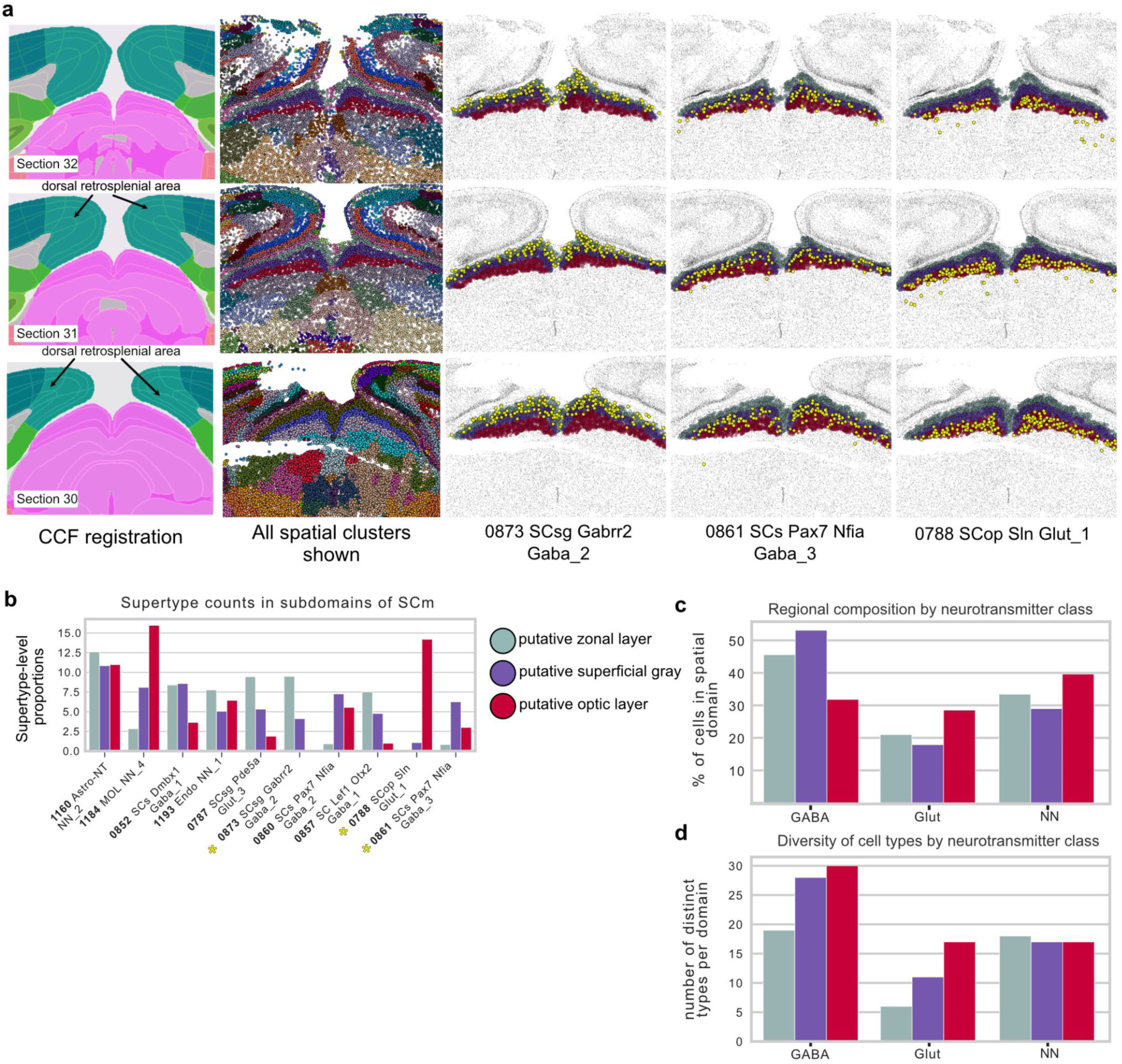
Comparison of best fit spatial domains from CellTransformer with layers of the superior colliculus, sensory related area. (**a.**) Sequential tissue sections (32, 31, 30, from anterior to posterior) showing in first column CCF registration and borders of relevant areas. Second column: all cells in field of view, colored by spatial domain from CellTransformer. Third column: only visualizing cells inside our putative matches for the zonal, superficial gray, and optic layers in the superior colliculus. The 0879 SCsg Pde5a Glut_1 cell type (supertype-level) in yellow. Fourth column: same as third but visualizing the 0865 SCs Pax7 Nfia Gaba_3 cell type. Fifth column: same as third and fourth but visualizing the 0882 SCop Sln Glut type. (**b.**) Bar chart of cell type abundance (as a percentage) for top ten most abundant types across the putative subregions. Cell types visualized in (**a.**) are marked with a yellow asterisk. (**c.**) Bar chart of per-region proportions of GABA-ergic and glutamatergic neurons and non-neuronal types. (**d.**) Bar chart of the number of distinct cell types at supertype level of the ABC-MWB taxonomy per domain.

**Supplementary Figure 10.**
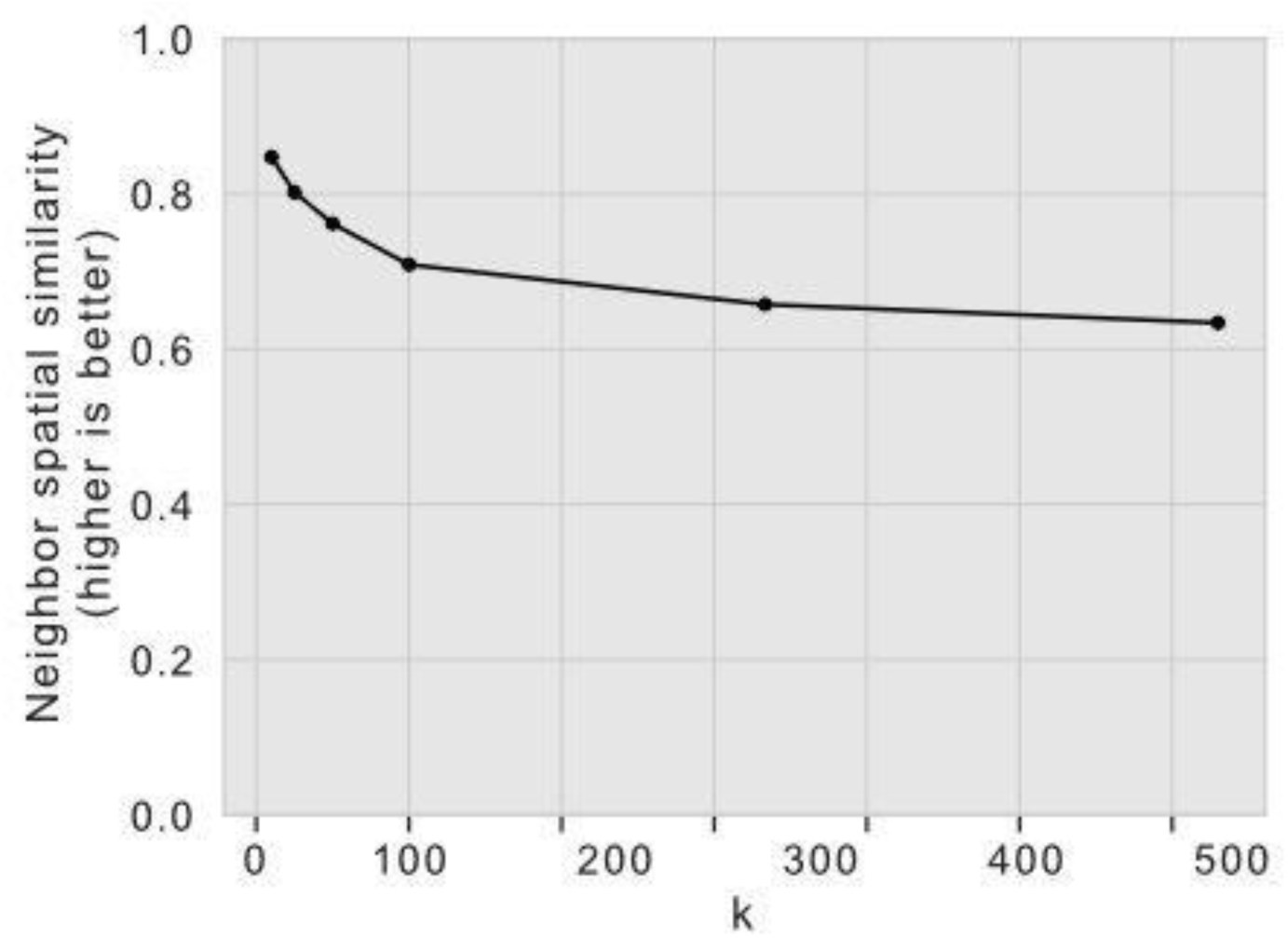
Results of quantitative comparison of CCF regions and CellTransformer regions at equivalent number of regions using the Zhuang 1-4 datasets^11^. Spatial smoothness of spatial clusters as measured using a nearest-neighbors approach, computed by clustering the concatenated latent variables for neighborhoods in the Zhuang lab datasets.

**Supplementary Figure 11.**
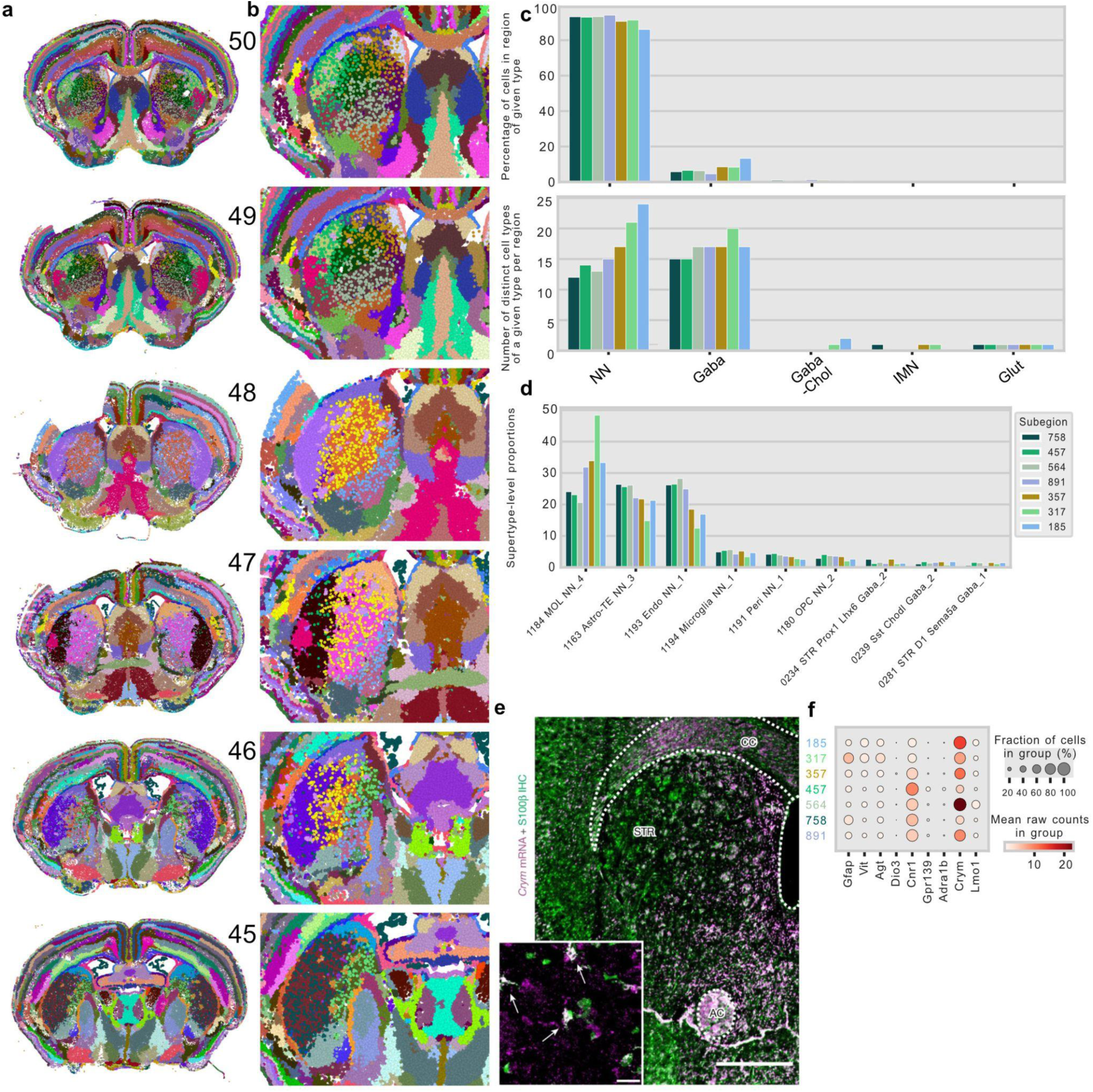
Representative images of spatial clustering from CellTransformer models with *k*=1300 identified using the Allen 1 dataset^1^. (**a.**) Sequential tissue sections (50 is most anterior) showing smoothness of spatial domains across and within tissue sections as well as consistent appearance of an irregular spatial pattern inside caudoputamen. (**b.**) Zoom in on the striatum for the same tissue sections. (**c.**) Plots showing percentage of cell types of different neurotransmitter for the non-uniform spatial clusters as well as the distribution of unique cell types of a given neurotransmitter type. (**d.**) Supertype-level counts in putative subpopulations of caudoputamen. (**e.**) Reproduction with permission of results from Ollivier et al. (2024). showing the distribution of *Crym* mRNA and its protein product (S100B), clearly identifying a medial population of *Crym*+ neurons which resembles the spatial pattern observed in clusters 758 and 457 (dorsoventral and Crym^-^). (**f.**) Dotplot of cell type expression proportions and mean counts per group (raw counts) in identified irregular caudoputamen areas.

**Supplementary Figure 12.**
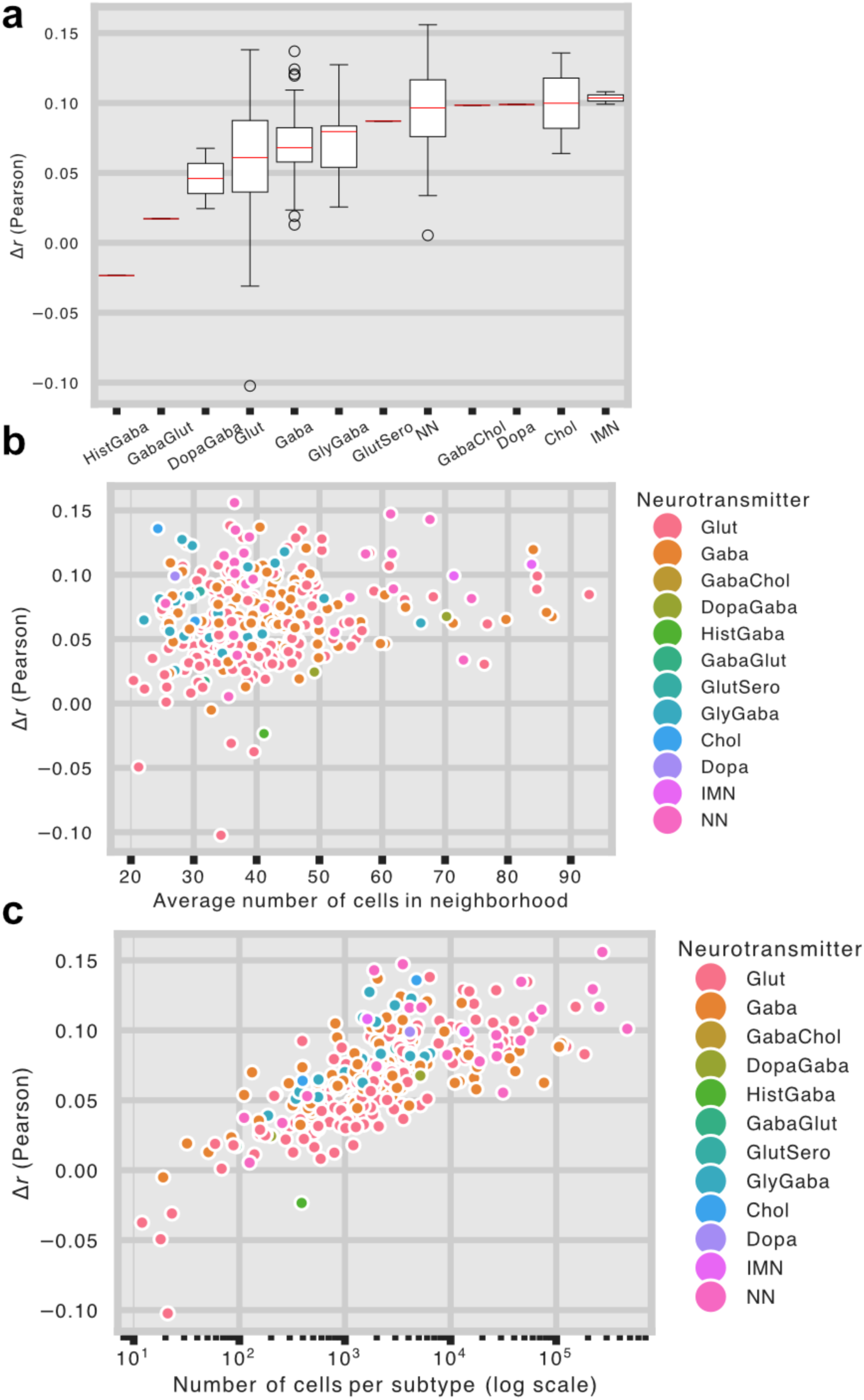
Quantification of improved prediction accuracy as a result of CellTransformer’s neighborhood-conditioned prediction. Results are over all cells in the Allen 1 dataset^1^. (**a.**) Change in Pearson correlation from per-cell type (at subclass level) average expression. Red lines show medians per distribution. (**b**.) Scatterplot of increase in average Pearson correlation per subclass level cell type against average neighborhood size for reference cells of that type. **c**. Scatterplot of increase in average Pearson correlation per subclass level cell type vs the number of cells of that type in log scale.

## References

1. Yao, Z. et al. A high-resolution transcriptomic and spatial atlas of cell types in the whole mouse brain. Nature 624, 317–332 (2023).

2. Consortium, T. Mic., et al. Functional connectomics spanning multiple areas of mouse visual cortex. 2021.07.28.454025 Preprint at 10.1101/2021.07.28.454025 (2023).

3. Chen, X. et al. Whole-cortex in situ sequencing reveals input-dependent area identity. Nature 1–10 (2024) doi:10.1038/s41586-024-07221-6.

4. Haviv, D. et al. The covariance environment defines cellular niches for spatial inference. Nat. Biotechnol. 1–12 (2024) doi:10.1038/s41587-024-02193-4.

5. Hu, J. et al. SpaGCN: Integrating gene expression, spatial location and histology to identify spatial domains and spatially variable genes by graph convolutional network. Nat. Methods 18, 1342–1351 (2021).

6. Zhou, X., Dong, K. & Zhang, S. Integrating spatial transcriptomics data across different conditions, technologies and developmental stages. Nat. Comput. Sci. 3, 894–906 (2023).

7. Townes, F. W. & Engelhardt, B. E. Nonnegative spatial factorization applied to spatial genomics. Nat. Methods 1–10 (2022) doi:10.1038/s41592-022-01687-w.

8. Birk, S. et al. Quantitative characterization of cell niches in spatial atlases. 2024.02.21.581428 Preprint at 10.1101/2024.02.21.581428 (2024).

9. Shi, H. et al. Spatial atlas of the mouse central nervous system at molecular resolution. Nature 622, 552–561 (2023).

10. Wang, Q. et al. The Allen Mouse Brain Common Coordinate Framework: A 3D Reference Atlas. Cell 181, 936–953.e20 (2020).

11. Zhang, M. et al. Molecularly defined and spatially resolved cell atlas of the whole mouse brain. Nature 624, 343–354 (2023).

12. Langlieb, J. et al. The molecular cytoarchitecture of the adult mouse brain. Nature 624, 333–342 (2023).

13. Veličković, P., et al. Graph Attention Networks. Preprint at 10.48550/arXiv.1710.10903 (2018).

14. Fischer, D. S., Schaar, A. C. & Theis, F. J. Modeling intercellular communication in tissues using spatial graphs of cells. Nat. Biotechnol. 41, 332–336 (2023).

15. Yao, Z. et al. A taxonomy of transcriptomic cell types across the isocortex and hippocampal formation. Cell 184, 3222–3241.e26 (2021).

16. Bakken, T. E. et al. Comparative cellular analysis of motor cortex in human, marmoset and mouse. Nature 598, 111–119 (2021).

17. Hintiryan, H. et al. The mouse cortico-striatal projectome. Nat. Neurosci. 19, 1100–1114 (2016).

18. Varrone, M., Tavernari, D., Santamaria-Martínez, A., Walsh, L. A. & Ciriello, G. CellCharter reveals spatial cell niches associated with tissue remodeling and cell plasticity. Nat. Genet. 56, 74–84 (2024).

19. Guo, T. et al. SPIRAL: integrating and aligning spatially resolved transcriptomics data across different experiments, conditions, and technologies. Genome Biol. 24, 241 (2023).

20. Zhang, X., Wang, X., Shivashankar, G. V. & Uhler, C. Graph-based autoencoder integrates spatial transcriptomics with chromatin images and identifies joint biomarkers for Alzheimer’s disease. Nat. Commun. 13, 7480 (2022).

21. Dong, K. & Zhang, S. Deciphering spatial domains from spatially resolved transcriptomics with an adaptive graph attention auto-encoder. Nat. Commun. 13, 1739 (2022).

22. Spatially informed clustering, integration, and deconvolution of spatial transcriptomics with GraphST | Nature Communications. https://www.nature.com/articles/s41467-023-36796-3.

23. Wu, S. et al. Stability-driven nonnegative matrix factorization to interpret spatial gene expression and build local gene networks. Proc. Natl. Acad. Sci. 113, 4290–4295 (2016).

24. Bienkowski, M. S. et al. Integration of gene expression and brain-wide connectivity reveals the multiscale organization of mouse hippocampal networks. Nat. Neurosci. 21, 1628–1643 (2018).

25. Cembrowski, M. S. et al. The subiculum is a patchwork of discrete subregions. eLife 7, e37701 (2018).

26. Bienkowski, M. S. et al. Homologous laminar organization of the mouse and human subiculum. Sci. Rep. 11, 3729 (2021).

27. Ding, S.-L. et al. Distinct Transcriptomic Cell Types and Neural Circuits of the Subiculum and Prosubiculum along the Dorsal-Ventral Axis. Cell Rep. 31, 107648 (2020).

28. Cang, J., Chen, C., Li, C. & Liu, Y. Genetically defined neuron types underlying visuomotor transformation in the superior colliculus. Nat. Rev. Neurosci. 25, 726–739 (2024).

29. Inagaki, H. K. et al. A midbrain-thalamus-cortex circuit reorganizes cortical dynamics to initiate movement. Cell 185, 1065–1081.e23 (2022).

30. McElvain, L. E. et al. Specific populations of basal ganglia output neurons target distinct brain stem areas while collateralizing throughout the diencephalon. Neuron 109, 1721–1738.e4 (2021).

31. Benavidez, N. L. et al. Organization of the inputs and outputs of the mouse superior colliculus. Nat. Commun. 12, 4004 (2021).

32. Cable, D. M. et al. Robust decomposition of cell type mixtures in spatial transcriptomics. Nat. Biotechnol. 40, 517–526 (2022).

33. Brbić, M. et al. Annotation of spatially resolved single-cell data with STELLAR. Nat. Methods 19, 1411–1418 (2022).

34. Maskey, S., et al. Generalized Laplacian Positional Encoding for Graph Representation Learning. arXiv.org https://arxiv.org/abs/2210.15956v2 (2022).

35. Burgess, J., Nirschl, J. J., Zanellati, M.-C., Cohen, S. & Yeung, S. Learning orientation-invariant representations enables accurate and robust morphologic profiling of cells and organelles. 2022.12.08.519671 Preprint at 10.1101/2022.12.08.519671 (2022).

36. Connell, W., Khan, U. & Keiser, M. J. A single-cell gene expression language model. arXiv.org https://arxiv.org/abs/2210.14330v1 (2022).

37. Laboratory, I. B. et al. A Brain-Wide Map of Neural Activity during Complex Behaviour. 2023.07.04.547681 Preprint at 10.1101/2023.07.04.547681 (2023).

38. de Vries, S. E., Siegle, J. H. & Koch, C. Sharing neurophysiology data from the Allen Brain Observatory. eLife 12, e85550 (2023).

39. Oh, S. W. et al. A mesoscale connectome of the mouse brain. Nature 508, 207–214 (2014).

40. Hike, D. et al. High-resolution awake mouse fMRI at 14 Tesla. eLife 13, (2024).

41. Paszke, A., et al. PyTorch: An Imperative Style, High-Performance Deep Learning Library. Preprint at 10.48550/arXiv.1912.01703 (2019).

42. Array programming with NumPy | Nature. https://www.nature.com/articles/s41586-020-2649-2.

43. Pedregosa, F. et al. Scikit-learn: Machine Learning in Python. J. Mach. Learn. Res. 12, 2825–2830 (2011).

44. Virtanen, P. et al. SciPy 1.0: Fundamental Algorithms for Scientific Computing in Python. Nat. Methods 17, 261–272 (2020).

45. Wolf, F. A., Angerer, P. & Theis, F. J. SCANPY: large-scale single-cell gene expression data analysis. Genome Biol. 19, 15 (2018).

46. Raschka, S., Patterson, J. & Nolet, C. Machine Learning in Python: Main developments and technology trends in data science, machine learning, and artificial intelligence. ArXiv Prepr. ArXi*v200204803* (2020).

47. Hunter, J. D. Matplotlib: A 2D graphics environment. Comput. Sci. Eng. 9, 90–95 (2007).

48. Waskom, M. L. seaborn: statistical data visualization. J. Open Source Softw. 6, 3021 (2021).

49. Ollivier, M. et al. Crym-positive striatal astrocytes gate perseverative behaviour. Nature 627, 358–366 (2024).

